# PARP1 condensates differentially partition DNA repair proteins and enhance DNA ligation

**DOI:** 10.1101/2024.01.20.575817

**Authors:** Christopher Chin Sang, Gaelen Moore, Maria Tereshchenko, Michael L. Nosella, Hongshan Zhang, T. Reid Alderson, Morgan Dasovich, Anthony Leung, Ilya J. Finkelstein, Julie D. Forman-Kay, Hyun O. Lee

## Abstract

Poly(ADP-ribose) polymerase 1 (PARP1) is one of the first responders to DNA damage and plays crucial roles in recruiting DNA repair proteins through its activity – poly(ADP-ribosyl)ation (PARylation). The enrichment of DNA repair proteins at sites of DNA damage has been described as the formation of a biomolecular condensate. However, it is not understood how PARP1 and PARylation contribute to the formation and organization of DNA repair condensates. Using recombinant human PARP1 *in vitro*, we find that PARP1 readily forms viscous biomolecular condensates in a DNA-dependent manner and that this depends on its three zinc finger (ZnF) domains. PARylation enhances PARP1 condensation in a PAR chain-length dependent manner and increases the internal dynamics of PARP1 condensates. DNA and single-strand break repair proteins XRCC1, LigIII, Polβ, and FUS partition in PARP1 condensates, although in different patterns. While Polβ and FUS are both homogeneously mixed within PARP1 condensates, FUS enrichment is greatly enhanced upon PARylation whereas Polβ partitioning is not. XRCC1 and LigIII display an inhomogeneous organization within PARP1 condensates; their enrichment in these multiphase condensates is enhanced by PARylation. Functionally, PARP1 condensates concentrate short DNA fragments and facilitate compaction of long DNA and bridge DNA ends. Furthermore, the presence of PARP1 condensates significantly promotes DNA ligation upon PARylation. These findings provide insight into how PARP1 condensation and PARylation regulate the assembly and biochemical activities in DNA repair foci, which may inform on how PARPs function in other PAR-driven condensates.

## INTRODUCTION

Cells are frequently exposed to DNA damaging agents such as reactive oxygen species or ionizing radiation that are detrimental to genome integrity. One of the first responders to DNA damage is poly(ADP-ribose) polymerase 1 (PARP1), the most abundant member of the human PARP family of proteins (1, 2) that covalently modify biomolecules with a monomer or polymer of ADP-ribose (3–5). PARP1 is allosterically activated (6) upon binding to single- and double-strand breaks in DNA (7, 8) and generates poly(ADP-ribose) or PAR on itself (9, 10) and proteins in the vicinity of the DNA lesion, such as histones (9, 11). This localized PARylation leads to the recruitment of other proteins to the site of damage (12, 13), including chromatin remodelers (14–16) and DNA repair proteins (17, 18). Accordingly, PARP1 and its activity are crucial in multiple single- and double-strand break repair pathways (19). Yet, how PARP1 influences the organization of damaged DNA and its target proteins and contributes to subsequent repair reactions is not well understood.

The enrichment of DNA repair proteins at DNA lesions, called DNA repair foci, has recently been described as biomolecular condensates (4, 20–22). Biomolecular condensates are non-membrane bound organelles that concentrate certain biomolecules and exclude others. Associative interactions between the molecules compensate for the entropic cost of condensation, leading to their demixing from the surrounding milieu as distinct phases in a process called phase separation (23–26). They are thought to regulate multiple biological processes including ribosome biogenesis (27, 28), stress responses (29, 30), and signal transduction (31, 32) by concentrating specific components and thus influencing biochemical reaction rates (24, 33). Post-translational modifications drastically alter the assembly, composition, and material properties of condensates by modulating interactions between biomolecules, as shown for phosphorylation (34–36), arginine methylation (34, 37–39), and O-linked-N-acetylglucosaminylation (40, 41). The rapid and extensive recruitment and autoPARylation of PARP1 in response to DNA damage raises the possibility that it serves as a seed to nucleate DNA repair condensate formation.

Among the proteins recruited to DNA lesions by PARylation are enzymes involved in single-strand break repair (SSBR), a well-characterized pathway in which PARP1 acts as the primary sensor of single-strand breaks, including single strand nicks. PARP1 activation at lesions leads to recruitment of SSBR proteins, including a scaffold protein XRCC1 that interacts with DNA, PARP1, and PAR (48–52) to bring together the proteins that will repair the break such as DNA polymerase β (Polβ) and DNA ligase III (LigIII) (44–47). Polβ and LigIII also interact with PARP1 and PAR (42, 43) that likely contributes to their recruitment. Condensate-forming proteins such as FUS-EWS-TAF15 (FET) family proteins also localize to DNA damage sites in a PARP1-activity dependent manner (4, 53, 54) and play important roles in SSBR (55). Whether and how PARP1 and its target proteins form DNA repair condensates and how their organization around damaged DNA is influenced by PARylation are unknown.

Here, we report that PARP1 forms condensates in a DNA-dependent manner. PARP1 auto-PARylation enhances its condensation and differentially promotes the partitioning of single-strand break repair proteins, FUS, Polꞵ, LigIII and XRCC1, within PARP1 condensates. Interestingly, PARP1 condensates concentrate short DNA, promotes compaction of long DNA, and bridge DNA ends. Upon PARylation, PARP1 condensates further enrich XRCC1-LigIII and enhance DNA ligation efficiency. Our findings support a model in which PARP1 nucleates condensates that selectively enrich and organize SSBR proteins at sites of DNA damage, promoting efficient ligation of DNA single-strand breaks following PARylation.

## RESULTS

### PARP1 forms condensates in DNA-dependent manner

Human PARP1 (UniProt ID: P09874) comprises three regions – a zinc finger (ZnF) region, automodification domain (AD), and catalytic region (CAT; **Fig. 1A**) – containing several folded domains (Refs: (8, 56, 57) and PDB-IDs: 2COK, 2CR9) interspersed with regions of disorder that are 20-50 amino-acids long (**Supp. Fig. 1A**). Three algorithms that predict the likelihood of phase separation based on either the sequence properties of intrinsically disordered protein regions or on sequence similarity to known RNA granule components differed dramatically in their estimates of whether PARP1 would undergo phase separation (**Supp. Table 1**). To test this experimentally, we examined the presence or absence of phase separation in solutions of recombinant mCherry-tagged human PARP1 *in vitro* (**Supp. Fig. 1B**) using fluorescence microscopy. mCherry-PARP1 concentrations were varied over a range containing the estimated cellular concentration of 1–2 μM measured in HeLa cells (57). mCherry-PARP1 did not form condensates on its own at any of the tested protein and salt concentrations (**Fig. 1B** and **Supp. Fig 1C**). However, the addition of a damaged DNA substrate – consisting of three oligonucleotides annealed to form a 60-nt triplex structure with a central nick and two blunt ends (triplex DNA; **Supp. Table 2**) – triggered the formation of micron-sized mCherry-PARP1 condensates. The presence of these condensates coincided with increasing PARP1 concentrations and at lower salt concentrations (**Fig. 1B** and **1C**).

**Figure 1.**
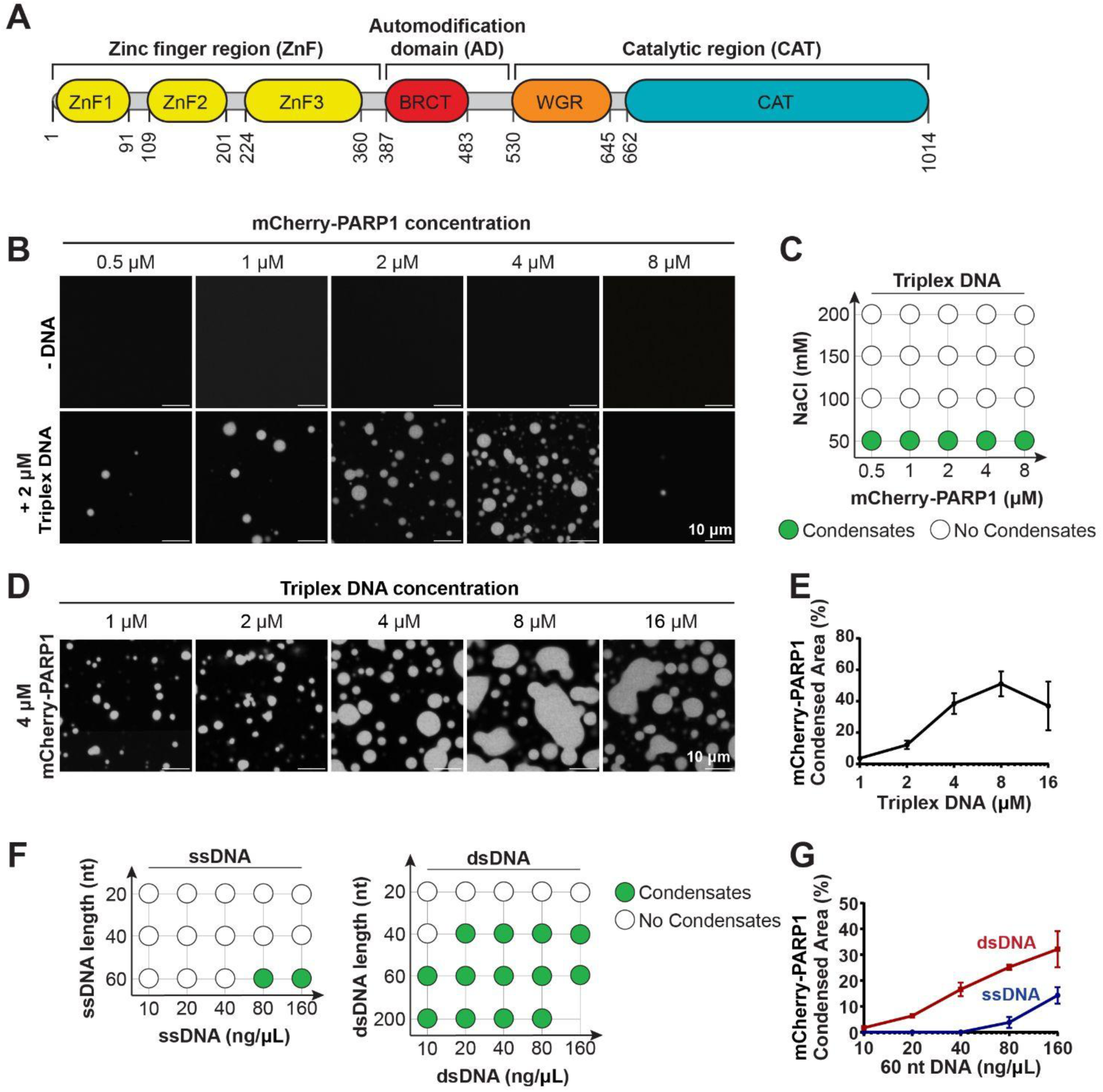
PARP1 forms condensates in a DNA-dependent manner. **A.** Domain architecture of PARP1. ZnF, zinc finger domain; BRCT, BRCA1 C-terminus domain; WGR, tryptophan-glycine-arginine region; CAT, catalytic domain. **B.** Fluorescence micrographs of condensates formed by recombinant mCherry-PARP1 in the presence or absence of DNA at the indicated concentrations in PARylation Buffer (20 mM Tris pH 7.5, 50 mM NaCl, 7.5 mM MgCl_2_, 1 mM DTT). **C.** Condensate formation by mCherry-PARP1 in PARylation Buffer in the presence of DNA and absence of NAD^+^. **D.** Fluorescence micrographs of 4 µM mCherry-PARP1 in PARylation Buffer with the indicated concentration of triplex DNA without NAD^+^. **E.** Quantifications of the surface area covered by mCherry-PARP1 condensates in D. Error bars indicate the standard error of the mean (SEM). **F.** Left: Condensate formation by 4 µM mCherry-PARP1 with the indicated ssDNA length and concentrations in PARylation Buffer in the absence of NAD^+^. Right: Condensate formation by 4 µM mCherry-PARP1 with the indicated dsDNA length and concentrations in PARylation Buffer in the absence of NAD^+^. **G.** Quantifications of the surface area covered by mCherry-PARP1 condensates in Supp. Fig 1F. Error bars indicate the SEM. All experiments were repeated in at least 3 independent replicates.

To further characterize the effects of DNA on PARP1 condensation, we quantified mCherry-PARP1 condensation in the presence of DNA of differing lengths and strandedness by fluorescence microscopy. We found that the nicked triplex DNA promoted mCherry-PARP1 condensation in a concentration-dependent manner (**Fig. 1D**, quantified in **Fig. 1E**). Increasing the concentration ratio of PARP1 to DNA led to a reduction in PARP1 condensation (**Fig. 1B**, quantified in **Supp. Fig. 1D**). Similarly, at higher DNA to PARP1 ratios, PARP1 condensates did not form as readily (**Fig. 1D and E**), suggesting reentrant phase behavior (58). In addition, PARP1 condensation was enhanced by increased concentration of a 25-nt oligonucleotide containing a single nick (nicked dumbbell DNA; **Supp. Table 2**) (**Supp. Fig. 1E**) and single-stranded and double-stranded oligonucleotides (ssDNA and dsDNA, respectively) (**Supp. Fig. 1F)**. Overall, dsDNA triggered mCherry-PARP1 condensation more readily compared to ssDNA, driving condensation at shorter lengths (**Supp. Fig. 1F,** depicted as phase diagrams in **Fig. 1F**, and quantified in **Fig. 1G**). Longer dsDNA also promoted PARP1 condensation at higher salt concentrations compared to a shorter double-stranded DNA, triplex DNA (**Supp. Fig. 1G** compared to **Fig. 1C**). Taken together, our data indicate that PARP1 undergoes condensation in a DNA-dependent manner and that DNA length and strandedness strongly influence this process. Based on the estimated DNA footprint of PARP1 on DNA being 14 nucleotides flanking a single-strand break (59), we conclude a DNA fragment longer than 20 nucleotides establishes the multivalency needed for PARP1 condensation.

### AutoPARylation enhances the formation and internal dynamics of PARP1 condensates

Upon binding to damaged DNA, PARP1 is allosterically activated and PARylates both itself and vicinal proteins. To examine whether PARP1 autoPARylation influences its condensation, we activated the enzyme by mixing it with the nicked triplex DNA and the substrate for PARylation, the ADP-ribose donor nicotinamide adenine dinucleotide (NAD^+^) (6, 8, 60–62). This resulted in near-complete autoPARylation of PARP1 within 15 minutes in a DNA- and NAD^+^-dependent manner, as observed by its reduced electrophoretic mobility (**Fig. 2A)** and positive reactivity on an anti-PAR immunoblot (**Supp. Fig 2A**), consistent with previous reports (6, 8, 60). PAR chain lengths were analyzed following cleavage and precipitation via chemoenzymatic labelling of terminal ADP-ribose moieties with Cy3-dATP and visualization on a urea-PAGE gel (103). PAR chains in our reaction ranged from 2 to 100+ ADP-ribose moieties, with the most abundant chain-lengths ranging from ∼10–60-mer (**Supp Fig. 2C**) at a total concentration of 355 µM (**Supp. Fig. 2B**). This corresponds to an approximate concentration of 10 µM assuming a consistent length of 35 units. AutoPARylated mCherry-PARP1 formed larger condensates compared to unmodified mCherry-PARP1 condensates (**Fig. 2B**, quantified in **Supp. Fig. 2D**), while not significantly changing the conditions at which microscopically visible PARP1 condensation occurred in the presence of 2µM triplex DNA (**Supp. Fig. 2E** compared to **Fig. 1C**). The presence of free 16-mer PAR chains alone (PAR16, (63)), at concentrations comparable to those in a PARylation reaction, did not enhance PARP1 condensation (**Supp. Fig. 2F**). This suggests that PARP1 autoPARylation is more potent at enhancing PARP1 condensation than free PAR chains.

**Figure 2.**
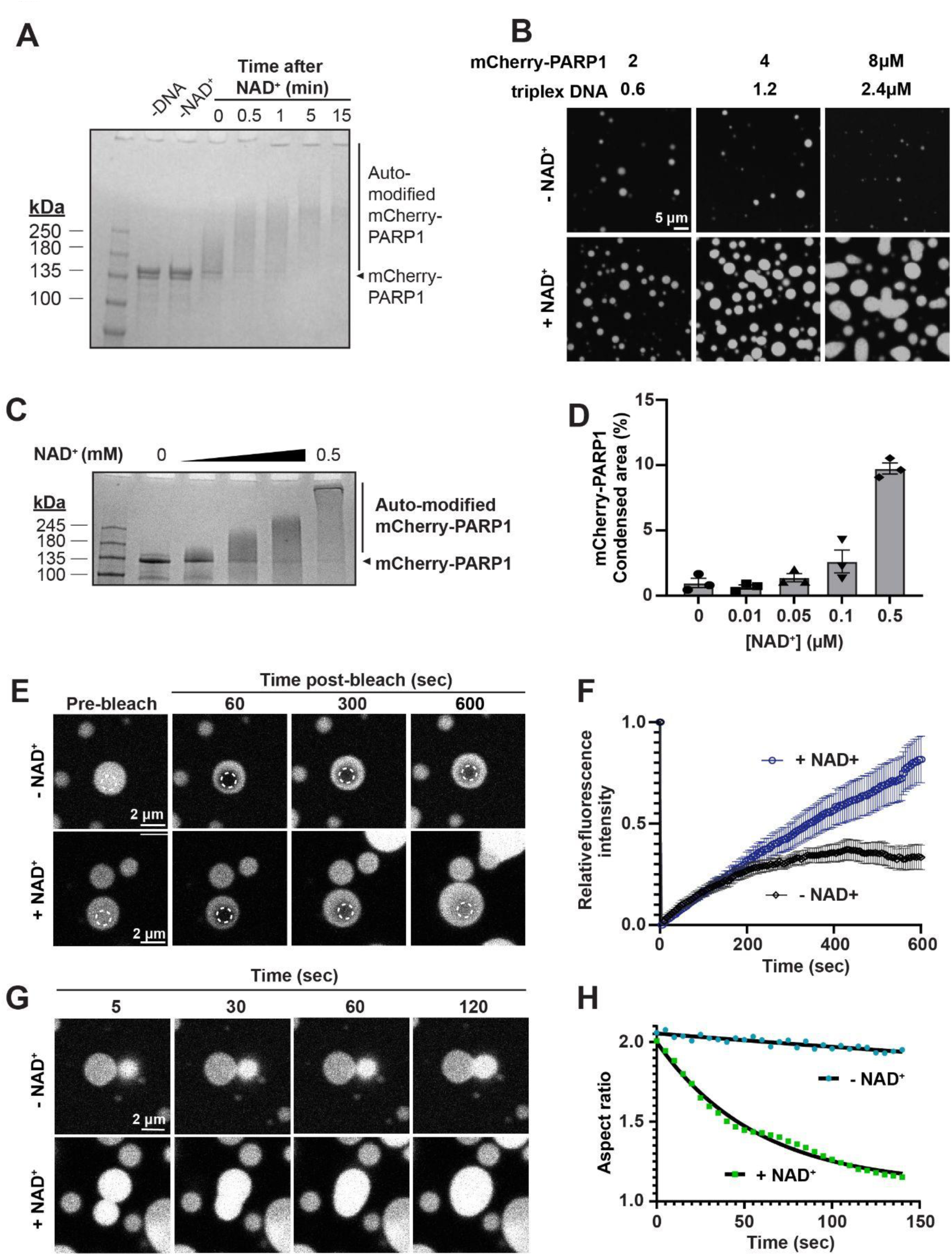
AutoPARylation enhances formation and internal dynamics of PARP1 condensates. **A**. SDS-PAGE of autoPARylation assay performed in 1 µM mCherry-PARP1 in PARylation Buffer (20 mM Tris pH 7.5, 50 mM NaCl, 7.5 mM MgCl_2_, 1 mM DTT, 0.3 µM dumbbell DNA). Time after addition of 0.5 mM NAD^+^ is indicated. Smearing and reduced electrophoretic mobility indicates autoPARylation. **B**. Fluorescence micrographs of the indicated concentration of mCherry-PARP1 in an autoPARylation reaction with the corresponding concentration of triplex DNA containing PARylation Buffer in the presence or absence of 0.5 mM NAD^+^. **C.** Gel-based autoPARylation assay performed with 1 µM mCherry-PARP1 in PARylation Buffer with increasing concentrations of NAD^+^ (0 mM, 0.01 mM, 0.05 mM, 0.1 mM, and 0.5 mM). **D**. Quantifications of the surface area covered by mCherry-PARP1 condensates in Supp. Figure 1H. Error bars indicate the standard error of the mean (SEM). **E**. Fluorescence recovery after photobleaching (FRAP) micrographs of 4 µM mCherry-PARP1 in PARylation Buffer with and without NAD^+^. Condensates were photobleached in the region indicated by the dotted circles, and fluorescence recovery was monitored over time; n = 20 from three independent experiments. **F**. FRAP quantifications represented in E. **G**. Fluorescence micrographs showing fusion of mCherry-PARP1 condensates with and without NAD^+^. Condensates of 4 µM mCherry-PARP1 were formed in PARylation Buffer with 0.5 mM NAD^+^. Time from initiation of fusion events is indicated. **H**. Plot of droplet aspect ratio over time for fusion events shown in G. All experiments were repeated in at least 3 independent replicates unless specified otherwise.

To determine how PAR chain length affects PARP1 phase separation, we limited the length of the chains on PARP1 by modulating the concentration of NAD^+^ or by titrating a competitive PARP1/2 inhibitor, ABT-888 (64), into the autoPARylation reaction mixture. Reducing the NAD^+^ concentration or increasing the ABT-888 concentration reduced autoPARylation of mCherry-PARP1 as observed by gel electrophoresis (**Fig. 2C** and **Supp. Fig. 2G**). Reducing the NAD^+^ concentration also corresponded with reduced condensate size when visualized by fluorescence microscopy (**Fig. 2D**, corresponding images in **Supp. Fig. 2H**). Increasing amounts of ABT-888 also decreased PARP1 condensate size (**Supp. Fig. 2I**). Together, these data show that autoPARylation enhances PARP1 condensation, which is proportional to the length of the PAR chains.

UnPARylated PARP1 condensates triggered by DNA were spherical, suggesting that their morphologies were influenced by surface tension as commonly observed in liquids. However, fluorescence recovery after photobleaching (FRAP) of a region within unPARylated mCherry-tagged PARP1 condensates was limited, plateauing at ∼25% normalized intensity, implying a mobile fraction of ∼25% (**Fig. 2E** and **2F, -NAD^+^**). This indicates that there is minimal internal rearrangement and exchange between the condensate and the surrounding environment, as often seen in highly viscous condensates. Consistent with this, adjacent condensates of unPARylated mCherry-PARP1 did not readily fuse or relax into spherical structures (**Fig. 2G** and **2H, -NAD^+^**). In contrast, the fluorescence of autoPARylated mCherry-PARP1 condensates recovered steadily, reaching close to complete recovery levels over time (**Fig. 2E** and **2F, +NAD^+^**). In addition, adjacent autoPARylated mCherry-PARP1 condensates fused readily, relaxing into a spherical shape within 2 minutes of touching (**Fig. 2G** and **2H, +NAD^+^**). A similar trend was observed using dumbbell DNA, suggesting that the type of DNA substrate does not influence this behavior (data not shown). Together, these data indicate that autoPARylation increases the mobility of molecules within PARP1 condensates.

### The ZnF region is required for PARP1 condensation

Given the importance of DNA for PARP1 condensation, we asked if the zinc finger region (ZnF), which binds DNA, is sufficient and necessary for this process. We generated two ZnF truncation mutants, PARP1-ΔZnF, which includes all regions of PARP1 except for the three ZnF domains, and PARP1-ZnF, which only contains the three ZnFs and excludes all other regions (**Fig. 3A** and **Supp.** Fig. 3A). We analyzed their phase separation alone or together at various protein concentrations in the absence or presence of triplex DNA and NAD^+^. In the absence of DNA, neither of the truncation mutants (PARP1-ΔZnF and PARP1-ZnF) condensed at concentrations as high as 32 μM (**Supp. Fig. 3B and 3D**). Upon the addition of DNA, PARP1-ZnF readily formed condensates at concentrations as low as 8 μM (**Fig. 3B** and **3C**), whereas PARP1-ΔZnF was unable to form condensates at any tested concentration (**Fig. 3C** and **Supp. Fig. 3E**). Deletion of either the ZnF region (PARP1-ΔZnF) or the remainder of the protein (PARP1-ZnF) abolished the autoPARylation activity of PARP1 (**Fig. 3F**), consistent with previous reports (8, 60, 65), and accordingly showed no NAD^+^-dependent change in condensation of the truncations (**Fig. 3B** and **Supp. Fig 3E**). These findings indicate that the interactions between the ZnF region and DNA are essential for PARP1 condensation and may be sufficient to drive the process at higher concentrations.

**Figure 3.**
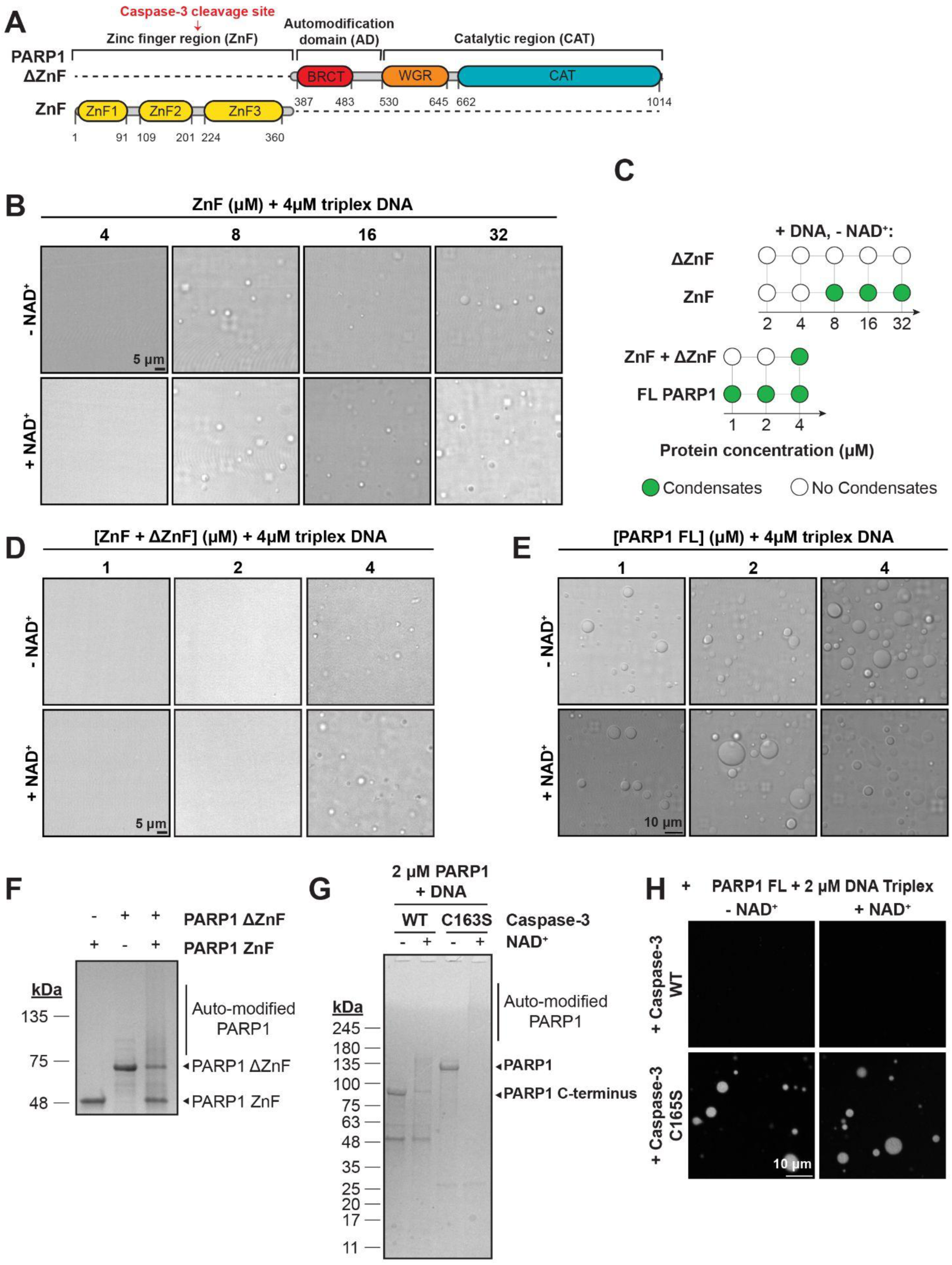
The ZnF region is important for PARP1 condensation. **A**. Domain architecture of truncated PARP1 proteins. **B**. DIC micrographs of ZnF PARP1 protein with 4 µM triplex DNA in PARylation Buffer (20 mM Tris pH 7.5, 50 mM NaCl, 7.5 mM MgCl2, 1 mM DTT). **C**. Condensate formation by truncated and full-length PARP1 proteins at the indicated concentrations in PARylation Buffer. **D**. DIC micrographs of truncated PARP1 proteins with or without 4 µM triplex DNA and their deleted domains (as indicated) in an autoPARylation reaction in the presence of 0.5 mM NAD^+^. **E.** DIC micrographs of full-length PARP1 proteins with 2 µM triplex DNA in an autoPARylation reaction with or without 0.5 mM NAD^+^. **F.** Gel-based autoPARylation assay of 1 µM truncated PARP1 proteins and their deleted domains in PARylation buffer in the presence or absence of 0.5 mM NAD^+^, analyzed 10 min after NAD+ addition. The absence of a protein band or smearing in assays containing NAD^+^ indicates modification with PAR. **G**. Gel-based cleavage assay of mCherry-PARP1 with caspase-3 WT or catalytically inactive caspase-3 C163S. 1 µM caspase-3 WT or C163S was added to 2 µM mCherry-PARP1 in PARylation buffer and incubated for 30 min. After incubation, 0.5 mM NAD^+^ was added as indicated and reactions were analyzed 15 min post-reaction initiation. **H.** Fluorescence micrographs of 2 µM mCherry-PARP1 in PARylation Buffer with 2 µM triplex DNA, and with or without NAD^+^, pre-cleaved with 1 µM caspase-3 WT or catalytically inactive mutant C163S for 30 min. Caspase-3 was added to mCherry-PARP1 30 min prior to initiation of PARylation to ensure full PARP1 cleavage, and images were obtained 10 min after reaction initiation. All experiments were repeated in at least 3 independent replicates unless specified otherwise.

To examine whether the linkage between the ZnF region and the rest of the PARP1 is important for condensate formation, PARP1-ZnF was mixed with PARP1-ΔZnF. An equimolar mixture of the truncations restored condensation at concentrations as low as 4 μM, which is lower than the concentration needed for ZnF-only condensates (8 μM) but higher than the concentration at which full-length (FL)-PARP1 condenses (1 μM) in the presence of DNA (**Fig. 3D** and **3E**). Mixtures of PARP1 truncations did not form condensates in the absence of DNA at all concentrations tested, just like FL-PARP1 (**Supp. Fig. 3D, 3F, and 3G**). Interestingly, autoPARylation was restored when PARP1-ΔZnF and PARP1-ZnF were mixed at equimolar concentrations as low as 1 μM (**Fig. 3F**) but did not significantly enhance condensation of the mixture (**Fig. 3D**). This suggests that the interactions between the ZnF and ΔZnF contribute to PARP1 condensation.

Next, we investigated whether the linkage of three ZnF domains in the ZnF region is important for condensation. We treated PARP1 with either caspase-3 WT, which cleaves PARP1 between ZnF2 and ZnF3 during apoptosis (66), or caspase-3 C163S, a catalytically-dead mutant (**Fig. 3G**). Treating mCherry-PARP1 with caspase-3 WT abolished condensation, irrespective of autoPARylation, while treatment with caspase-3 C163S did not affect mCherry-PARP1 condensation (**Fig. 3H**). These results highlight that the three ZnFs in tandem are essential for PARP1 condensation. Together, the interaction of the ZnF region with the rest of the protein drives PARP1 condensation with DNA.

### PARP1 condenses long DNAs and bridge DNA ends

To understand the interaction of PARP1 and damaged DNA within condensates, we mixed mCherry-PARP1 with Cy5-labeled triplex DNA. In the absence of NAD^+^ (and thus of PARylation), Cy5-triplex DNA was enriched ∼20-fold in PARP1 condensates when compared to the concentration in the dilute phase (**Fig. 4A**, -NAD^+^), as determined by the fluorescence intensity measurements inside and outside the condensed phase. In the presence of NAD^+^ (and thus of PARylation), Cy5-triplex DNA was enriched ∼12-fold in PARP1 condensates compared to the dilute phase (**Fig. 4A**, +NAD^+^), which was less than pre-PARylation. Thus, PARP1 condensates significantly enrich damaged DNA with or without PARylation.

**Figure 4:**
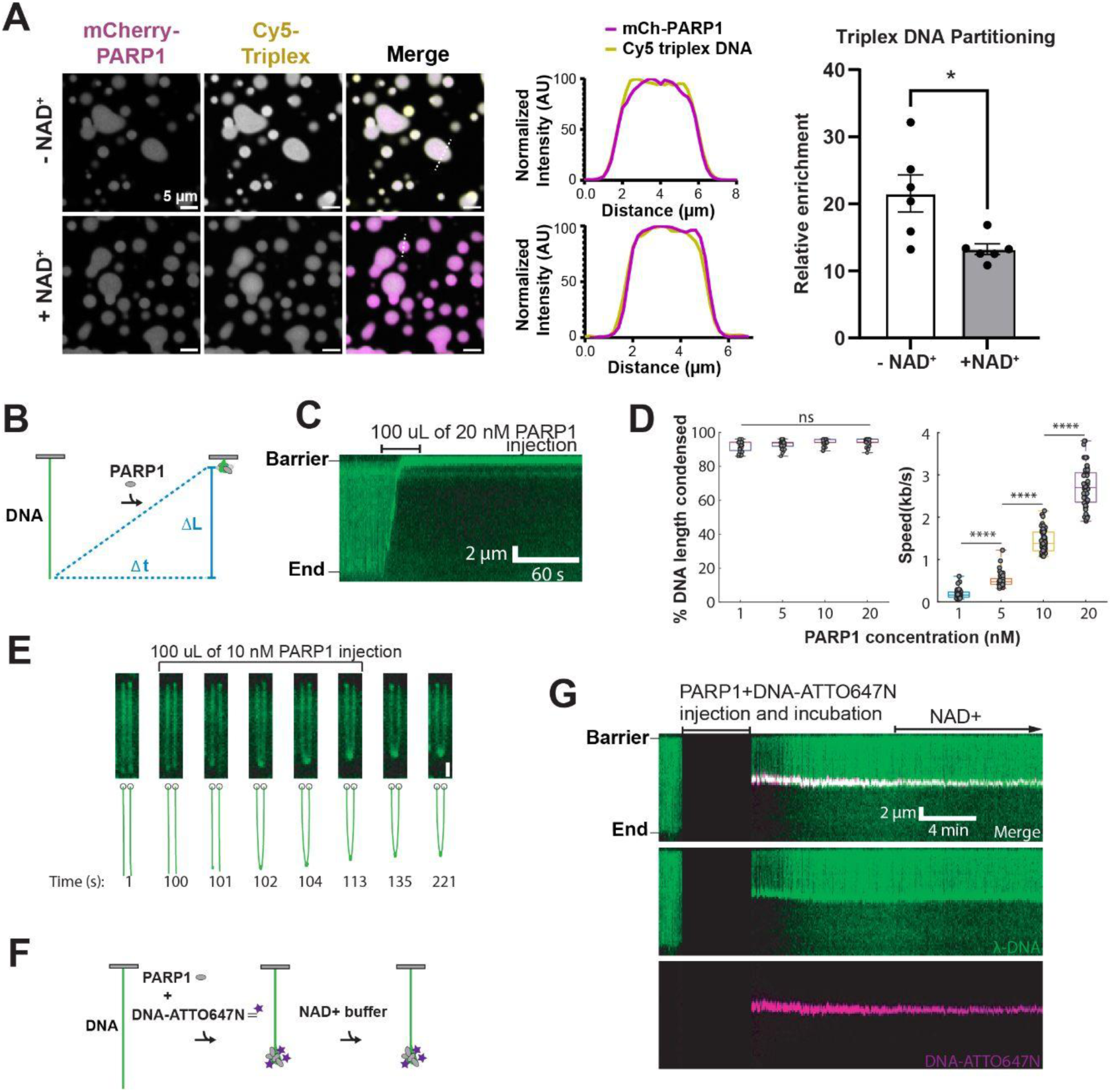
PARP1 condensates bridge broken DNA ends. **A.** Fluorescence micrographs of 4 µM mCherry-PARP1 and 4 µM Cy5-triplex DNA in PARylation Buffer (20 mM Tris pH 7.5, 50 mM NaCl, 7.5 mM MgCl_2_, 1 mM DTT) with and without NAD+. The fluorescence intensity measured across the diameter of a representative (dashed white line) condensate is plotted in the center. Quantifications of Cy5-triplex into mCherry-PARP1 condensates calculated as the fluorescence intensity in the condensed phase relative to the dilute phase; n = 6. P-value is obtained from a student t-test; * p < 0.05. **B.** Schematic of DNA condensation initiated by PARP1. The change in DNA length (ΔL) over a time (Δt) is used to calculate the mean DNA compaction rate (ΔL/Δt). **C.** Kymograph showing DNA (green) compaction after the injection of 100 μL of 20 nM PARP1. **D.** DNA compaction rates are dependent on the PARP1 concentration, with complete compaction observed at all tested concentrations. N>48 DNA molecules for all concentrations. p-values are obtained from a two-tailed t-test: ****: p < 0.0001, ns: not significant. **E.** Frames from a movie showing the end-to-end bridging of two DNA molecules following the injection of 100 μL of 10 nM PARP1. Scale bar: 2 μm. **F.** Schematic of the 60 bp dsDNA capture experiment. **G.** Kymographs illustrating the capture of the 60 bp DNA by PARP1. 100 μL containing 5 nM PARP1 and 20 nM ATTO647N-oligo DNA was injected into the flowcell with flow turned off. The flow was resumed and reaction was imaged after all components were incubated for five minutes. Chasing this reaction with a buffer containing 500 μM NAD+ did not release the captured oligo. All experiments were repeated in at least 3 independent replicates unless specified otherwise.

To understand how PARP1 interacts with and influences longer DNA molecules, we examined PARP1 in a single molecule DNA curtains assay (67). In this assay, a 48.5 kb λ-DNA substrate is tethered to the passivated surface of a microfluidic flowcell via a biotin streptavidin linkage. The DNA was stained with the intercalating dye SYTOX Orange and wild type PARP1 was injected into the flowcell at the indicated concentration (**Fig. 4B**). After PARP1 injection, DNA molecules compacted in a PARP1 concentration-dependent manner (**Fig. 4C** and **4D** and **Supp. Fig. 4A**). The DNA compacted by 92.2±0.4% to 94.4±0.3% when 1-20 nM of PARP1 continuously entered the flowcell (N>48 for all conditions). The rate of DNA compaction was dependent on the concentration of PARP1, with compaction speeds of 0.18±0.02 kb/s for 1 nM PARP1, 0.51±0.02 kb/s for 5 nM PARP1, 1.43±0.03 kb/s for 10 nM PARP1, and 2.73±0.07 kb/s for 20 nM PARP1. The compacted DNA remained stable after the addition of 500 mM NAD^+^ (**Supp. Fig. 4B**). This suggests that PARP1 clusters in DNA curtain assays can also concentrate long DNA, capturing and condensing it in *cis* with or without PARylation.

Interestingly, injection of a sub-saturating 100 μL of 10 nM PARP1 into the flowcell led to end-to-end bridging between two adjacent DNA molecules, indicating that PARP1 can hold two DNA ends in *trans* (**Fig. 4E** and **Supp. Movie 1**). The bridged DNA ends withstood applied forces of approximately 0.17±0.02 pN, which was estimated using the Worm-Like-Chain model (68, 69). To further characterize this phenomenon, we designed a double stranded oligonucleotide capture assay. Here, a mixture of 5 nM PARP1 and 20 nM ATTO647N-labeled 60 bp dsDNA was introduced into the flowcell and incubated for 5 minutes with buffer flow turned off (**Fig. 4F**). Subsequently, the flow was resumed and images were acquired. All PARP1-DNA clusters were co-localized with DNA-ATTO647N, indicating that PARP1 clusters could capture non-complementary DNA in *trans* and bridge interactions between the oligonucleotide and long DNA substrate (**Fig. 4G**). As before, these clusters could not be resolved by 500 nM of NAD^+^ (**Supp. Fig. 4C**). Together these data support a possibility that PARP1 condensates condense and bridge long DNA.

### PARP1 condensation and activity enhance nicked DNA ligation

A possible function of PARP1 condensation is to organize and concentrate DNA repair enzymes and their substrates. To test this, we investigated how PARP1 phase separation contributes to the organization of several SSBR proteins: the DNA ligase LigIII, its obligatory binding partner XRCC1, the DNA polymerase Polβ, and FUS (**Supp.** Fig. 5A). When mixed with mCherry-PARP1, AlexaFluor488-labeled XRCC1 and LigIII were heterogeneously distributed within PARP1 condensates individually (**Fig. 5A** and **Supp. Fig 5B**, **-NAD^+^**) and together (**Fig. 5D, -NAD**^+^). The heterogeneous partitioning of XRCC1 and LigIII did not change upon PARylation (**Fig. 5A** and **5D**, and **Supp. Fig 5B, +NAD^+^**). PARylation did enhance the overall enrichment of LigIII and XRCC1 in PARP1 condensates by approximately 2-fold (**Fig. 5A, 5B**, and **5C**). Interestingly, when LigIII and XRCC1 were added together, PARP1 demixed from the XRCC1/LigIII complex and DNA, such that areas enriched in mCherry-PARP1 were depleted in triplex DNA and LigIII whereas LigIII-rich areas strongly partitioned damaged DNA while depleting mCherry-PARP1 (**Fig. 5D**). This behavior is reminiscent of multi-phase condensates in which two or more phases coexist within a condensate, such as in nucleoli (27, 70).

**Figure 5.**
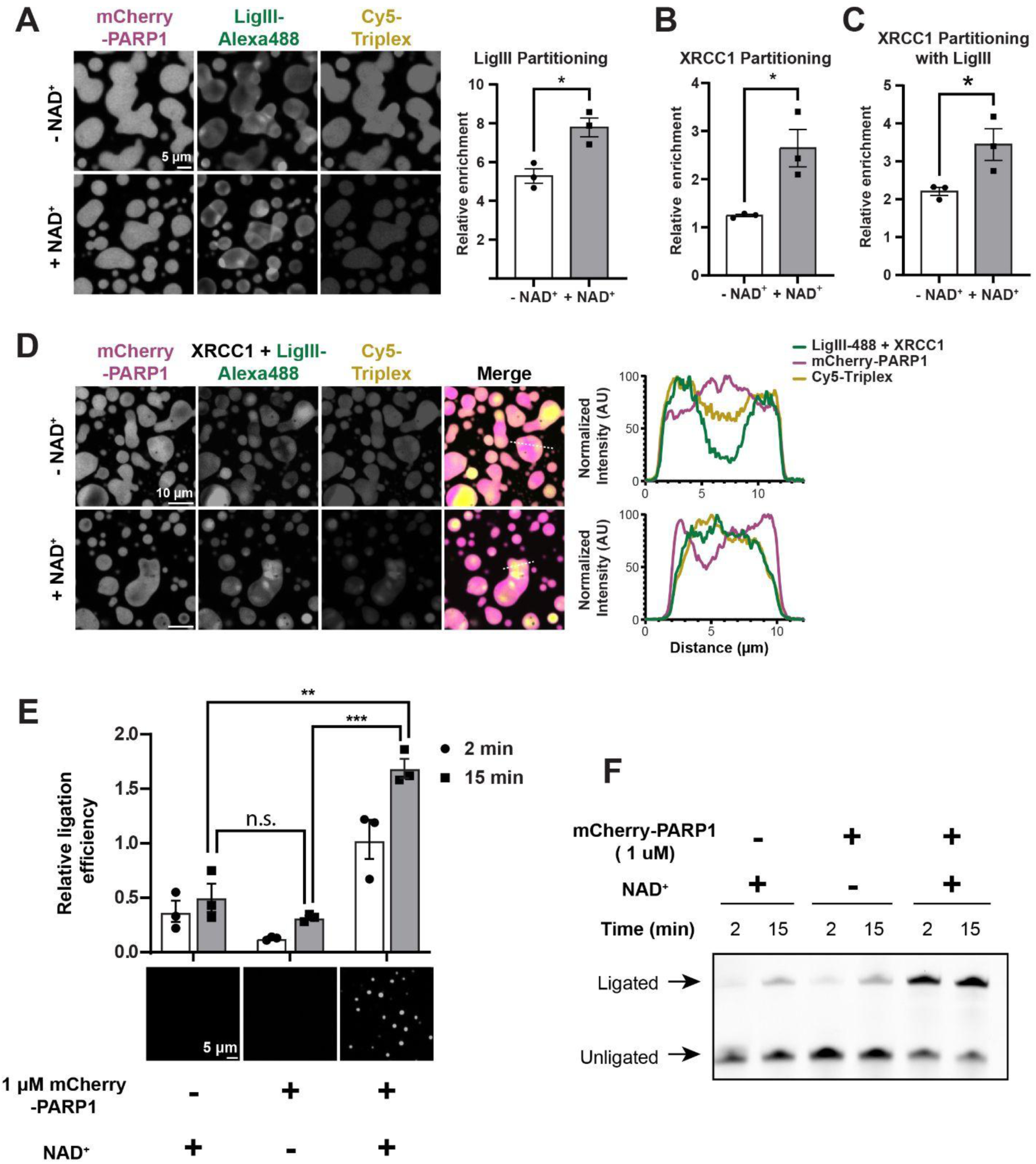
SSBR proteins partition together in condensates that enrich DNA. **A.** Left: Fluorescence micrographs of 4 µM mCherry-PARP1, 4 µM triplex DNA and 1 µM LigIII (10% AlexaFluor488-labeled LigIII) in PARylation Buffer (20 mM Tris pH 7.5, 50 mM NaCl, 7.5 mM MgCl_2_, 1 mM DTT) with NAD^+^. Right: Quantification of the partitioning of LigIII within condensates as calculated by the mean fluorescence intensity in the condensed phase relative to that of the dilute phase. **B.** Quantification of the partitioning of XRCC1 in Supp. Fig. 5B containing 4 µM mCherry-PARP1, 4 µM triplex DNA (10% Cy5-labeled triplex DNA) and 1 µM XRCC1 (10% AlexaFluor488-labeled XRCC1) in PARylation Buffer with and without NAD^+^. The enrichment of the XRCC1 within condensates as calculated by the mean fluorescence intensity in the condensed phase relative to that of the dilute phase is plotted. **C.** Quantification of the partitioning of XRCC1 in Supp. Fig. 5C (10 minutes) containing 4 µM mCherry-PARP1, 4 µM triplex DNA, 1 µM LigIII, and 1 µM XRCC1 (10% AlexaFluor488-labeled XRCC1) in PARylation Buffer with and without NAD^+^. The enrichment of the XRCC1 within condensates as calculated by the mean fluorescence intensity in the condensed phase relative to that of the dilute phase is plotted. **D.** Fluorescence micrographs of 4 µM mCherry-PARP1, 4 µM triplex DNA and 1 µM LigIII (10% AlexaFluor488-labeled LigIII) and 1 µM XRCC1 in PARylation Buffer with NAD^+^. The fluorescence intensity measured across the diameter of a representative condensate (white dashed line) is plotted on the right. **E.** Top: Gel-based ligation assay performed in PARylation Buffer with 100 nM LigIII, 100 nM XRCC1, 50 nM Cy5-triplex DNA and with or without 1 µM mCherry-PARP1 and 0.5 mM NAD^+^, analyzed 2 or 15 min after ATP addition. Quantifications of ligation efficiency calculated as the fluorescence intensity of Cy5-triplex DNA in the upper band (ligated) relative to the lower band (unligated), as shown in F. Bottom: Representative fluorescence micrographs of mCherry-PARP1 in the ligation reactions after ATP addition. **F.** Gel-based ligation assay performed in PARylation Buffer with 100 nM LigIII, 100 nM XRCC1, 50 nM Cy5-triplex DNA and with or without 1 µM mCherry-PARP1 and 0.5 mM NAD^+^ analyzed 2 or 15 min after ATP addition. The presence of a DNA band at higher molecular weight indicates ligation. Quantifications are shown in E. All experiments were repeated in at least 3 independent replicates unless specified otherwise. p-values are obtained from a student t-test: *** p < 0.001, ** p < 0.01, * p < 0.05, n.s.: not significant.

Another SSBR protein, AlexaFluor488-labeled Polβ, was evenly enriched in the PARP1 condensates (∼1.5 fold) (**Supp. Fig 5C, -NAD^+^**), and its partitioning in PARP1 condensates did not change with PARylation (**Supp. Fig 5C, +NAD^+^** and **5D**). FUS-GFP was also evenly enriched within PARP1 condensates (∼1.5 fold) (**Supp. Fig. 5E, -NAD^+^**) but its partitioning was significantly increased (∼4-fold) upon PARylation (**Supp. Fig 5E, +NAD^+^** and **5F**). Based on previous reports that XRCC1 and FUS strongly interact with PAR (48, 71), their increased partitioning to PARylated PARP1 condensates may be driven by PAR binding. Together, our data indicate that PARP1 condensates and PARylation differentially organize SSBR proteins, XRCC1, LigIII, Polβ, and FUS.

Next, we investigated how PARP1 condensation influences DNA repair efficiency by monitoring the ligation of a nicked DNA substrate in the presence of XRCC1, LigIII, and PARP1 with and without NAD^+^ over time. We found that the presence of PARP1, without condensate formation, could be slightly inhibitory to ligation (**Fig. 5E** and **5F**). Interestingly, in the presence of PARP1 condensates (triggered with NAD^+^), ligation increases nearly 3-fold (**Fig. 5E** and **5F**). Our data suggests that PARP1 condensates increase DNA ligation potentially by enriching LigIII, XRCC1, and damaged DNA upon PARylation.

## DISCUSSION

In this study, we examine the role of PARP1 and PARylation in the formation, organization, and function of nascent, multi-component DNA repair foci. We report that PARP1 readily forms viscous condensates in a manner dependent on the concentration of DNA and PAR, thus expanding the list of DNA repair proteins known to undergo phase separation. Modification of PARP1 by PARylation enhances formation of PARP1 condensates and has differing effects on the organization of DNA repair proteins within them. Functionally, PARP1 condensates concentrate short DNA, which correlates with PARP1 clusters compacting and bridging long DNA ends in a single molecule DNA curtains assay. Furthermore, the activity of PARP1 condensates enhances DNA ligation. Together, these findings suggest a model whereby autoPARylation of PARP1 seeds the formation of condensates at DNA lesions that support efficient DNA repair (**Fig. 6**).

**Figure 6.**
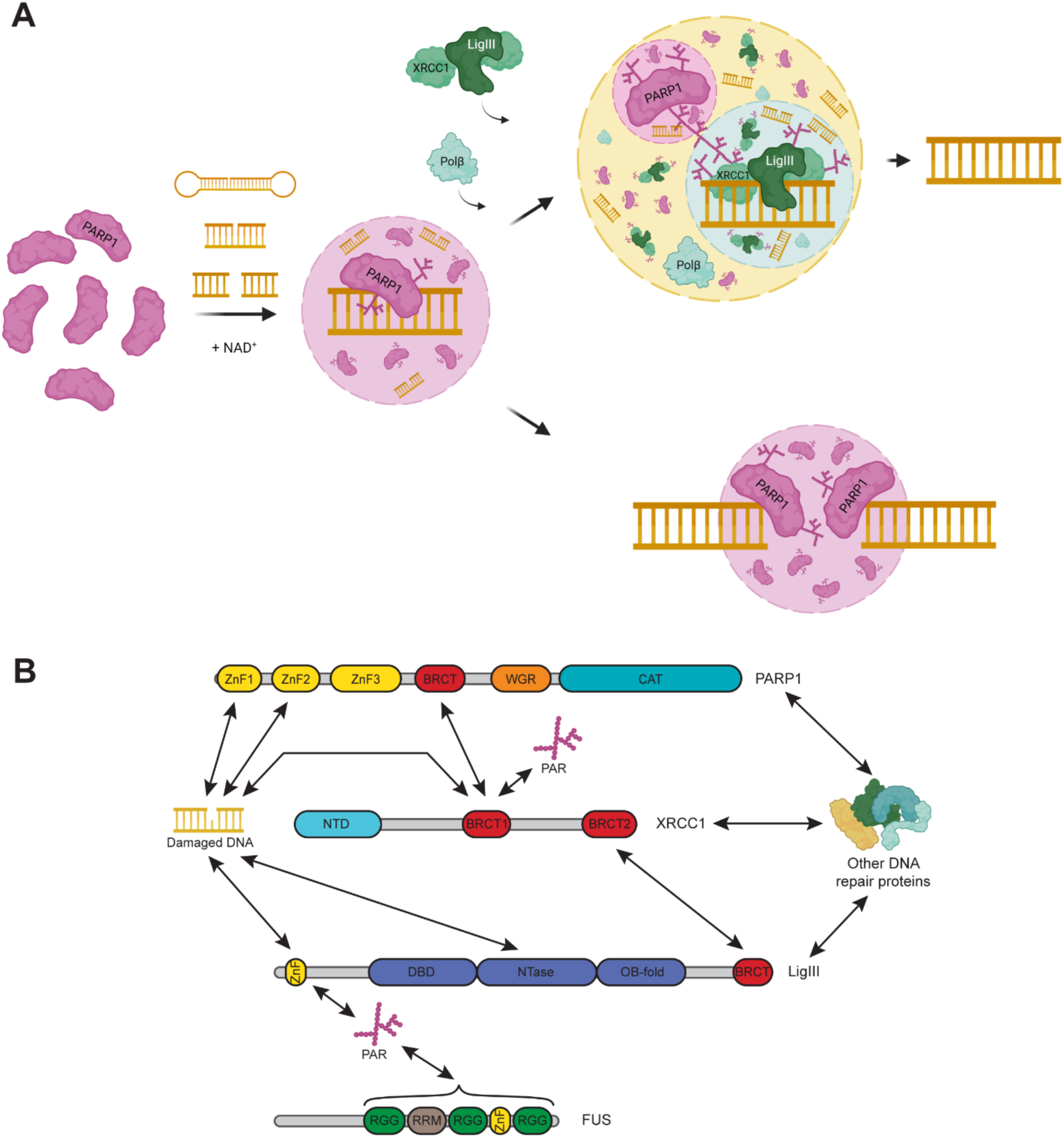
Model of PARP1 condensation in DNA repair. **A.** When monomeric PARP1 binds a DNA lesion (nicked dumbbell, nicked triplex, or double-strand break), it is allosterically activated and PARylates itself which triggers the formation of a PARP1 condensate that enriches damaged DNA. The role of PARP1 condensation varies according to the type of DNA lesion. The recruitment of single-strand break repair proteins XRCC1, LigIII, and Pol ꞵ to nicked DNA substrates leads to the formation of multiphase condensates and, in combination with PARylation, the enhancement of DNA ligation efficiency. Alternatively, PARP1 condensates can compact and bridge broken DNA ends together. **B.** Schematic showing the reported interactions of PARP1, XRCC1, LigIII, and FUS with each other, PAR, and DNA. The figures were created with BioRender.

PARP1 is an interesting example of a predominantly folded protein that undergoes condensation. Unlike many other phase-separating proteins such as the FUS-like RNA-binding proteins, which have disordered regions hundreds of residues long (72), PARP1 has only short ∼10–20 residue disordered regions interspersed between multiple highly structured interaction domains. PARP1 does not form condensates on its own but requires the presence of DNA to do so (**Fig. 1**). This is similar to other nucleic acid-binding proteins, such as G3BP1 (73) and VRN1 (74), whose condensation requires the presence of RNA and DNA, respectively. Nucleic acids promote condensation by adding to the multivalency of the system through interactions with each other as well as bringing associated proteins into closer proximity. In our system, multivalency of the system is further increased by the three tandem ZnF domains in PARP1 (**Fig. 3**) and by the PAR chains, which are nucleic acid-like polymers of up to 200 ADP-ribose moieties (1, 2, 63). Our finding that ZnF domains are essential for PARP1 condensation is interesting as other regions of PARP1 have been shown to also interact with DNA (104). The strength of interactions between DNA and PARP1 regions may contribute to these differences. Supporting this notion, we find that the first two ZnF domains that have been shown to directly sense and bind single strand nicks with high affinity (106–108) are necessary for PARP1 condensation. Interestingly, we find that further increasing DNA concentration dissolves PARP1 condensation as PARP1 undergoes a reentrant phase transition (**Fig. 1**). RNA-binding proteins, including FUS, hnRNPA1, TDP-43, have been shown to display similar reentrant phase behavior (75), as the protein-protein interactions are diluted by increased protein-RNA interactions. A similar phenomenon likely contributes to PARP1 reentrant behavior. In cells, there are limited pools of naked DNA available with most bound tightly by histones. Thus, PARP1 condensation would occur predominantly at nucleosome-depleted DNA breaks and potentially transcription sites.

Our finding that auto-PARylation enhances the formation of PARP1 condensates (**Fig. 2**) provides further evidence to support the hypothesis that PARylated proteins act as seeds for condensate formation in cells (4, 71, 76). Although free PAR chains enhance *in vitro* condensate formation of various RNA-binding proteins, such as the FET proteins, hnRNPA1, and TDP-43 (77–80), free PAR chains did not enhance PARP1 condensation, suggesting that PAR chains must be conjugated to facilitate the process. Recent findings suggest that two other PARP family members, PARP5a’s catalytic domain and full-length PARP7, form condensates *in vitro* upon ADP-ribosylation (80, 81) and this may be a generalizable behavior of PARPs. Moreover, our work shows that PAR chain length is crucial in this process, with short PAR chains promoting condensation much less effectively than long chains. Of note, PARylation increased internal mobility of PARP1 condensates in our study, which is in contrast to other studies that showed free PAR reduced the internal dynamics of condensates (78). Given the increased dynamics of PARP1 condensates upon PARylation, it is tempting to speculate that higher PAR concentration may even dissolve PARP1 condensates, similar to PARP1 reentrant behavior with high concentrations of DNA. Investigating other properties of PAR, such as branching (63), should provide further insights into how PARylation regulates PARP1 condensates.

We show that PARP1 condensates concentrate ∼60 nt-long DNA. Similarly, clusters of PARP1 that form at much lower concentrations than the micron-sized PARP1 condensates can compact long DNA of over 48,000 base pairs. This is consistent with results obtained with atomic force microscopy that PARP1 can cluster DNA (82). Our data that PARP1 clusters can bridge long DNA ends also raise the possibility that PARP1 condensates may bridge, and potentially protect, naked and broken DNA ends. Protecting and bridging broken DNA ends would be a crucial first step to efficient DNA repair in multiple repair pathways that often progress into double-stranded breaks (83). A recent study reported that a condensate forming protein, FUS, can compact and bridge DNA ends (84), similar to PARP1. It would be interesting to examine whether this is a common feature of many or specific proteins that phase separate. While PARP1 autoPARylation did slightly reduce the partitioning of shorter DNA fragments, it did not block enrichment of DNA fragments or compaction of longer DNA. This was surprising to us, given the model that PARylation causes PARP1 to release from DNA (6, 85) and causes decondensation of nucleosome arrays (86, 87). In line with this, we recently reported that PARylation in the presence of HPF1 blocks oligomerization of nucleosome core particles (105). HPF1 directs PARylation to occur mainly on serine residues, which is a major PARylation site observed in cells (109), and reduces overall PARylation *in vitro*. Thus, the presence of nucleosomes and/or general reduction of PARylation may contribute to the differences we observe in PARP1-DNA interactions following PARylation compared to previous studies. In addition, there are likely differences in how DNA partitions into PARP1 condensates compared to how PARP1 monomer interacts with DNA. We also report that PARylation alters PARP1 condensate composition by influencing the partitioning of SSBR proteins. PARylation promotes XRCC1 and FUS enrichment in PARP1 condensates. LigIII, XRCC1 and FUS possess PAR-binding domains (ZnF, BRCT1, and RGG repeats and RRMs, respectively) (**Fig. 6B**) (42, 48–50), which would enable them to be further enriched upon PARylation. On the other hand, PARylation did not influence the enrichment of Polβ (**Supp. Fig. 5D**). Further investigation will elucidate how competing affinities for DNA, PAR and other biomolecules lead to the spectrum of enrichment phenotypes at DNA repair foci. PARylation-dependent enrichment of SSBR proteins serves as the first instance, to our knowledge, of compositional control of biomolecular condensates by PARylation and may have implications in other condensates regulated by this post-translational modification, such as stress granules (3, 78, 79, 88) and the nucleolus (70, 89).

DNA ligation was enhanced in the presence of PARP1 condensates (**Fig. 5**). In this assay, PARylation was used to trigger PARP1 condensation. As such, PARP1 condensation and PARylation may both contribute to promoting DNA ligation. How might this work? PARylation further enriches XRCC1/LigIII to PARP1 condensates, and their increased concentration may facilitate increased DNA ligation. PARylation also increases the dynamics of PARP1 condensates, which may contribute to increased ligation by facilitating rapid exchange of proteins and DNA. A similar phenomenon was observed in SPOP/DAXX condensates where increased condensate dynamics correlated with increased catalytic activity of an E3 ubiquitin ligase, CRL3 (91). It is worth noting that DNA ligation was enhanced by 3-fold in the presence of PARP1/LigIII/XRCC1 condensates, which account for less than 5% of the total reaction mixture. This result was different from our recent finding that PARylation in the presence of HPF1 did not influence ligation of 601 DNA (105). An intriguing possibility is that increased overall PARylation or PARylation of LigIII enhances its condensate partitioning and/or activity. Our data also offer explanations for how PARP1 can interact with LigIII and XRCC1 through domain-domain interactions (42–47) (**Fig. 6B**), yet can compete with LigIII for binding to damaged DNA (90). We demonstrate that PARP1 and LigIII/XRCC1 form distinct, yet coexisting phases and damaged DNA prefers the LigIII/XRCC1 phase. The presence of XRCC1 was important for increased DNA partitioning with the LigIII/XRCC1 phase, suggesting that XRCC1 interactions with DNA or PARP1 or LigIII drive differential partitioning of DNA.

An open question is how PARP1 condensates dissolve. PAR formation at sites of damage is rapidly counteracted by PAR-degrading enzymes, including poly(ADP-ribose) glycohydrolase (PARG) and ADP-ribosyl hydrolase 3 (ARH3) (1). PARG, which preferentially cleaves ADP-ribose from the ends of PAR chains (92), arrives at DNA damage sites with similar kinetics as PARP1 (17). PARylation also triggers the recruitment of numerous repair proteins that also bind DNA at the damage sites (17, 18). Based on our data that PAR chain-length and DNA binding regulate PARP1 condensation, it seems plausible that the combination of PARG activity and arrival of other proteins that compete for DNA binding leads to PARP1 condensate dissolution over time. In addition, our evidence that cleavage of PARP1 by the cell death protease caspase-3 reduces its condensation suggests that during apoptosis, PARP1 condensate dissolution may be promoted by caspase-3 activity.

In summary, our results demonstrate that interactions between PARP1 and naked or damaged DNA leads to the formation of biomolecular condensates that enrich DNA and DNA repair proteins with functional consequences in holding together broken DNA ends and enhancing DNA ligation. Our findings support a model in which PARP1 phase separation and activity form a seed that enhances condensation of SSBR proteins into sub-compartments that preferentially concentrate different biomolecules. These findings shed insight into how PARP1 may facilitate SSBR protein recruitment and repair reactions. Future studies in cells would be valuable to understand how these effects at the molecular level contribute to DNA repair in a complex environment of the nucleus.

## MATERIALS AND METHODS

### Cloning and construct generation

Gibson assembly (New England Biolabs) was used to generate all constructs. cDNAs encoding full-length, wild-type *Homo sapiens PARP1*, *PARG*, *LIG3* (splice isoform alpha, containing C-terminal BRCT domain), and *XRCC1* were obtained from NIH Mammalian Gene Collection (MGC). DNA fragments containing the full-length coding sequences of each protein (PARP1 1-1014, PARG 1-976, XRCC1 1-633, LIG3 43-1009, excluding mitochondrial targeting sequence) were amplified by polymerase chain reaction (PCR) using Kapa HiFi Hotstart ReadyMix (Roche) and inserted into a pET-SUMO expression vector containing an N-terminal hexahistidine-SUMO tag (Invitrogen). PARP1 truncations (PARP1-ZnF, residues 1-378; PARP1-ΔZnF, residues 379-1014) were generated by PCR from PARP1 cDNA and inserted into a pET-SUMO expression vector. The DNA for full-length, wild-type human caspase-3 was synthesized by GenScript (Piscataway, NJ, USA), codon-optimized for expression in *E. coli*, and subsequently cloned into a pET-SUMO expression vector with an additional C-terminal hexahistidine tag cloned in using QuikChange (Agilent).

### Protein expression and purification

#### PARP1, PARP1-ZnF, PARP1-ΔZnF, PARG, XRCC1, Polβ

Full-length human PARP1 and truncation constructs were purified as described previously (8, 93, 94), with several modifications. Briefly, pET-SUMO expression vectors with His-SUMO-tagged PARP1 constructs were transformed into *E. coli* BL21 (DE3) RIPL cells and grown in LB with kanamycin and chloramphenicol. After overnight growth for 16 hr at 37 °C and 200 rpm, overnight cultures were used to inoculate large-scale cultures at a starting OD of ∼0.2. Construct expression was induced at an OD of ∼1.0 with 0.2 mM isopropyl β-D-1-thiogalactopyranoside (IPTG) and 0.1 mM zinc chloride and grown at 16 °C for 16-20 hr at 200 rpm. Cells were harvested by centrifugation and lysed in buffer containing 25 mM HEPES pH 7.5, 250 mM NaCl, 20 mM imidazole, 10% v/v glycerol, and 5 mM 2-mercaptoethanol with DNase and lysozyme by sonication (30% amplitude, 2s pulses/50% duty for 9 min total). Lysed cells were sedimented by centrifugation at 20,000 RCF at 4 °C for 30 min and the lysate was subsequently loaded onto pre-equilibrated Ni-NTA agarose resin (Cytiva) in a gravity column at room temperature. The resin was washed with 5 column volumes of lysis buffer and protein of interest was eluted using a step-wise elution in lysis buffer containing 50 mM, 100 mM, and 250 mM imidazole. Eluted protein was subjected to His-SUMO tag cleavage with ULP1 protease (purified in-house) in dialysis buffer containing 25 mM HEPES pH 7.5, 250 mM NaCl, 5 mM 2-mercaptoethanol overnight at 4 °C. Cleaved protein was subsequently separated from the His-SUMO tag and protease by re-loading onto Ni-NTA resin and collecting the flow-through in lysis buffer. The flow-through was then concentrated in an Amicon centrifugal unit (EMD-Millipore) with the appropriate molecular weight cutoff and purified on an AKTA FPLC system (Amersham Biosciences Co.) using a Superdex 200 16/600 HiLoad column (Cytiva) pre-equilibrated in 25 mM Tris pH 7.5, 1 M NaCl, 0.5 mM EDTA, 1 mM tris(2- carboxyethyl)phosphine (TCEP) at 4 °C. Fractions containing the protein of interest were pooled and concentrated in an Amicon centrifugal unit (EMD-Millipore) with the appropriate molecular weight cutoff and purified on an AKTA FPLC system (Amersham Biosciences Co.) using a pre-packed 4-mL source Q anion exchange resin (Cytiva) pre-equilibrated in 25 mM HEPES pH 7.5, 100 mM NaCl, 0.5 mM EDTA, and 5 mM 2-mercaptoethanol at 4 °C. Fractions containing protein of interest were pooled and dialyzed into 25 mM Tris pH 7.5, 250 mM NaCl, 0.5 mM EDTA, 5 mM 2-mercaptoethanol and concentrated. Purity was assessed by SDS-PAGE. PARG, XRCC1, and Polβ were purified using the same protocol, but bacterial cultures were not supplemented with ZnCl_2_.

#### LigIII

LigIII-alpha expression and purification was modified from previous literature (95). Briefly, a pET-SUMO expression vector with His-SUMO-tagged LigIII-alpha was transformed into *E. coli* BL21 (DE3) RIPL cells and grown in LB with kanamycin and chloramphenicol. After growth of overnight cultures for 16 hr at 37 °C and 200 rpm, overnight cultures were used to inoculate large-scale cultures at a starting OD of ∼0.2. Construct expression was induced at an OD of ∼0.6 with 0.3 mM isopropyl β-D-1-thiogalactopyranoside (IPTG) and grown at 16 °C for 16-20 hr at 200 rpm. Cells were harvested by centrifugation and lysed in buffer containing 50 mM Tris pH 7.5, 300 mM NaCl, 10% glycerol, 20 mM imidazole, 1 mM benzamidine, 2 mM 2- mercaptoethanol with DNase and lysozyme by sonication (30% amplitude, 2s pulses/50% duty for 9 min total). Lysed cells were sedimented by centrifugation at 20,000 RCF at 4 °C for 30 min and the lysate was subsequently loaded onto pre-equilibrated Ni-NTA agarose resin (Qiagen) at 4 °C. The resin was washed with 5 column volumes of lysis buffer and protein of interest was eluted using a step-wise elution with lysis buffer containing 100 mM, 200 mM, and 600 mM imidazole. Eluted protein was subjected to His-SUMO tag cleavage with ULP1 protease in dialysis buffer containing 50 mM Tris pH 7.5, 300 mM NaCl, 2 mM 2-mercaptoethanol overnight at 4C. Cleaved protein was subsequently separated from the His-SUMO tag and protease by re-loading onto Ni-NTA resin and collecting the flow-through in lysis buffer. The flow-through was then loaded onto a HiTrap Blue HP column (Cytiva) in 50 mM Tris pH 7.5, 300 mM NaCl, 10% glycerol, 1 mM benzamidine, 2 mM 2-mercaptoethanol. The column was washed with the same buffer and eluted with a step-wise salt gradient (500 mM NaCl, 1 M NaCl, 2 M NaCl). The elution fractions were then pooled and concentrated in an Amicon centrifugal unit (EMD-Millipore) with a 100 kDa MWCO membrane and purified on an AKTA FPLC system (Amersham Biosciences Co.) using a Superdex 200 pg 16/600 HiLoad column (Cytiva) pre-equilibrated in 25 mM Tris pH 7.5, 500 mM NaCl, 0.5 mM EDTA, 1 mM tris(2-carboxyethyl)phosphine (TCEP) at 4 °C. Fractions containing the protein of interest were concentrated and purity was assessed by SDS-PAGE.

#### Caspase-3

Full-length caspase-3 was expressed and purified as outlined in previous literature (96), with additional details included below for clarity. Caspase-3 initially exists in its zymogenic form (procaspase-3) and requires proteolytic cleavage for complete activation. Procaspase-3 is activated via cleavage at D9 and D28 in the prodomain, which subsequently dissociates, and D175 in the intersubunit linker to yield the p20 and p10 subunits, which then form a dimer of dimers (p20_2_p10_2_) (97). Because procaspase-3 is activated during overexpression in *E. coli*, any affinity tag that precedes the prodomain will dissociate from mature caspase-3 upon activation. For this reason, a C-terminal hexahistidine tag was cloned into the caspase-3 expression vector. The caspase-3 expression vector was transformed into *E. coli* BL21(DE3) competent cells, which were grown in LB medium at 37 °C with 30 μg/mL kanamycin and shaking at 220 rpm. When the optical density reached an OD of 0.6 units, the temperature was lowered to 16 °C and isopropyl β-D-1-thiogalactopyranoside (IPTG) was added to a final concentration of 0.2 mM. Caspase-3 expression proceeded for 18 hours at 16 °C and the cells were harvested by centrifugation and stored at -80°C until use. The cell pellet was resuspended in 25 mM HEPES, 150 mM NaCl, 0.5 mM 2-mercaptoethanol, and 30 mM imidazole at pH 8.0. Following lysis by sonication and clarification of the lysed material by centrifugation, the supernatant was loaded onto a 5 mL HisTrap HP column (Cytiva) in the same buffer listed above. Caspase-3 was eluted with 25 mM HEPES, 150 mM NaCl, 0.5 mM 2-mercaptoethanol, and 300 mM imidazole at pH 8.0. The eluted material was concentrated in an Amicon Ultra-15 3K MWCO concentrator and loaded onto a Hi-Load 16/600 Superdex 200 pg column (Cytiva) in buffer containing 25 mM HEPES, 50 mM NaCl, 5 mM tris(2- carboxyethyl)phosphine (TCEP) at pH 8.0. Fractions that contained caspase-3 were pooled, concentrated in an Amicon Ultra-15 3K MWCO concentrator and stored at -80 °C until use.

#### FUS-GFP

Full-length FUS tagged with a C-terminal GFP tag and an N-terminal His-MBP tag was overexpressed in BL21 (DE3) RIPL *E. coli* cells at 16 °C for 16 h after induction with 500 μM IPTG. Cells were harvested and lysed by sonication in lysis buffer (50 mM Tris-HCl, 500 mM NaCl, pH 8.0, 10 mM imidazole, 4 mM β-mercaptoethanol, 1 mM PMSF, and 0.1 mg/mL RNase A). The cell pellet was removed after centrifugation (16000 rpm, 4 °C, 1 h) and supernatant was loaded into a Ni-NTA agarose column (Qiagen). Protein was eluted in an elution buffer (50 mM Tris-HCl, 500 mM NaCl, pH 8.0, 100 mM imidazole and 4 mM β-mercaptoethanol). The His-MBP tag was cleaved using a His-tagged TEV protease during dialysis against FUS dialysis buffer (50 mM Tris-HCl pH 7.4, 1 M KCl, 10% glycerol, 4 mM β-mercaptoethanol) overnight at 4 °C. The mixture was loaded onto a Ni-NTA agarose column and FUS-GFP was collected in the flow-through fractions. The protein was further purified by gel filtration chromatography (HiLoad Superdex 200 pg 16/600; Cytiva), concentrated and equilibrated with storage buffer (50 mM Tris-HCl, 500 mM KCl, 2 mM DTT and 10% glycerol). Peak fractions were pooled and aliquoted in PCR tubes, flash-frozen in liquid nitrogen and stored at -80 °C.

#### Fluorescent protein labeling

1-10 μmol of protein of interest was dialyzed into 25 mM HEPES pH 7.5, 250 mM NaCl, 0.5 mM EDTA, 0.5 mM tris(2-carboxyethyl)phosphine (TCEP) overnight using 3.5kDa MWCO Slide-A-Lyzer dialysis cassettes (ThermoFisher Scientific). The protein was combined with 5x molar excess of Alexa Fluor 488 NHS-ester dye (ThermoFisher Scientific) dissolved in DMSO and incubated overnight at 4 °C in the dark. The reaction was quenched with 5x excess volume of 25 mM Tris pH 7.5, 250 mM NaCl, 0.5 mM EDTA, 5 mM 2-mercaptoethanol and desalted into the same buffer using a 5 mL HiTrap desalting column (GE Healthcare) using an AKTA FPLC system (Cytiva) at 4 °C. Fractions containing labeled protein were visualized by Alexa Fluor 488 fluorescence on a BioRad ChemiDoc MP system with Alexa Fluor 488 excitation/emission settings. Fractions containing labeled protein but no free dye were pooled and concentrated to desired concentration and stored at -80 °C.

#### *In vitro* PARylation assay

PARP1 at the indicated concentrations was combined with 0.3 μM nicked dumbbell DNA (**Supp. Table 2**) in PARylation Buffer (20 mM Tris pH 7.5, 50 mM NaCl, 7.5 mM MgCl_2_, and 1 mM DTT). 0.5 mM NAD^+^ was added to initiate the reaction and the reaction progression and/or phase separation were analyzed by SDS-PAGE, or microscopy at indicated timepoints. For experiments involving additional proteins (i.e. XRCC1-LigIII), these components were added to the reactions at the indicated concentrations prior to reaction initiation (unless indicated otherwise). For experiments involving labeled DNA oligonucleotides, Cy5-labeled triplex DNA (sequence found in **Supp. Table 2**) was added to the reaction in lieu of the dumbbell DNA at the indicated concentrations prior to reaction initiation. For experiments involving ABT-888 (Toronto Research Chemicals Inc), ABT-888 was added to reactions immediately prior to reaction initiation.

#### Western blot

Proteins were resolved onto a 4-20% Mini-PROTEAN TGX Precast gels (BioRad) and then transferred to a 0.2 μM nitrocellulose membrane (Biorad) in cold transfer buffer (25 mM tris base, 190 mM glycine and 20% methanol) for 1 hour at 90V (4℃). The nitrocellulose membrane was then blocked with blocking buffer (5% w/v skim milk powder, 0.05% v/v tween-20 in tris-buffered saline (TBST)) before its incubation with primary antibody (anti-PAR) overnight at 4℃. The membrane was then incubated in horseradish peroxidase (HRP)-linked secondary antibody (anti-mouse IgG) for 1 hour at room temperature and developed on a Biorad ChemiDoc MP imaging system using Luminata Crescendo Western HRP Substrate (Sigma, WBLUR0500).

#### Expression and purification of PARP5a (1093–1327)

PARP5a catalytic domain was expressed and purified as described (80).

#### Expression and purification of OAS1

OAS1 was expressed and purified as described (98).

### PAR synthesis and fractionation to PAR16

PAR was prepared essentially as described (98). PAR was synthesized enzymatically by PARP5 catalytic domain (0.1 mg/mL) with histones (2 mg/mL, Sigma #H9250) and NAD^+^ (20 mM) in 50 mM HEPES pH 7, 10 mM MgCl_2_, 1 mM DTT, 0.02% v/v NP-40 for 2 hrs at ambient temperature. PARylated proteins were precipitated with an equal volume of 20% w/v trichloroacetic acid (10% w/v final) and pelleted with 20,000 *g* for 30 min at 4 °C. Pellets were washed with 70% ethanol, then PAR was cleaved from the proteins with 0.5 M KOH, 50 mM EDTA for 1 hr at 60 °C. Sodium acetate pH 5.2 was added to a final concentration of 0.3 M, then ethanol was added to a final concentration of 70% v/v. PAR was precipitated for 1 hr at -80 °C, then pelleted with 20,000 *g* for 30 min at 4 °C. PAR pellets were washed with 70% ethanol, dried at 37 °C for 10 min, then resuspended in water. The mixture of PAR lengths was fractionated to PAR16 essentially as described (98). A preparative DNAPac PA100 column (22 x 250 mm) fractionated PAR into homogeneous polymers. PAR (up to 20 µmol ADPr) was loaded onto the column equilibrated with Dionex buffer A (25 mM Tris-HCl pH 9.0), and the concentration of Dionex buffer B (25 mM Tris-HCl pH 9.0 and 1M NaCl) in a 120-min method was set to elute as follows: 0 min (0% B), 6 min (0% B), 10 min (30% B), 60 min (40% B), 78 min (50% B), 108 min (56% B), 112 min (100% B), 114 min (100% B), 115 min (0% B), 120 min (0% B). Fractions containing PAR16 were combined, concentrated, and desalted into water with Amicon centrifugal filters (3k MWCO). [PAR16] was measured with a NanoDrop OneC using the equation: [PAR16] = *n* x A_260_ / 13,500 M^-1^ cm^-1^, where *n* is the number of adenines, in this case 16.

#### PAR chain length analysis of a PARP1 autoPARylation reaction

PAR from a PARP1 autoPARylation reaction was isolated, fluorescently labeled, and analyzed with PAGE essentially as described (98). AutoPARylated mCherry-PARP1 was acid precipitated, and PAR was isolated with base and ethanol precipitation as described above for PAR16 synthesis. The mixture of PAR lengths (∼250 µM ADPr) was labeled with Cy3-dATP (10 µM, Jena Bioscience #NU-835-CY3), poly(I:C) RNA (50 µg/mL, Invivogen #tlrl-picw), and OAS1 (1 µM) in 25 mM HEPES pH 7.5, 20 mM MgCl_2_, 2.5 mM DTT at 37 °C for 2 hrs. Labeling reactions were diluted with Formamide-EDTA loading buffer and separated with 12% urea-PAGE (National Diagnostics # EC-833) in an adjustable slab gel apparatus (VWR #CBASG-400) equipped with 28 cm plates. Cy3 signal was measured with a Licor-M. The image was exported and annotated with ImageStudio and Adobe Illustrator.

#### SDS-PAGE

Protein samples were quenched in 4x SDS-PAGE sample buffer (Biorad) to a final concentration of 1x. Samples were loaded onto 4-12% gradient gels (Biorad) and resolved for 25 min at 200V, and then stained with Coomassie blue (prepared in-house) and destained overnight. Gels were imaged on a BioRad GelDoc EZ system.

#### DIC microscopy of XRCC1, and LigIII condensates

PARylation reactions were set up as described above and transferred onto 35mm-diameter glass-bottom dishes (MatTek) and sealed with a glass coverslip to limit evaporation. Images were acquired on a Nikon Ti-2E microscope with an Andor Dragonfly 200 unit using a 40x oil immersion objective (DIC channel).

#### Fluorescence microscopy

PARylation reactions were set up as described above, transferred onto 35mm-diameter glass-bottom dishes (MatTek), and sealed with a glass coverslip to limit evaporation. Images were acquired on a Nikon Ti-2E microscope with an Andor Dragonfly 200 unit using a 60x oil immersion objective and 2048 x 2048-pixel resolution. Fluorescence was detected using a Zyla sCMOS camera after excitation with 521 nm, 600 nm, and 700 nm lasers. Images were processed and analyzed using Fiji. Photobleaching was performed using the LASX FRAP module on the Leica SP8 microscope. Circular regions of interest (ROI) of 1 μm x 1 μm size were positioned in the center of droplets (PARP1) or around the full droplet (XRCC1 and LigIII). Bleaching was performed with the 488 nm laser at 30% laser power for 1 repetition on zoom-in mode, and images were taken at 1.29s intervals. Fluorescence recovery over time was normalized to the intensity of images taken pre-bleach and plotted in GraphPad Prism. For condensate area analysis, the area covered by condensates was determined by setting an intensity threshold and dividing by the area of the whole image. For protein enrichment analysis, the average fluorescence intensity within condensates was divided by the average fluorescence intensity in the dilute phase.

#### Single-molecule fluorescence microscopy

Single-molecule fluorescent images were collected using a customized prism TIRF microscope (PMID:31780627). An inverted Nikon Ti-E microscope system was equipped with a motorized stage (Prio ProScan II H117). The flowcell was illuminated with a 488 nm laser (Coherent Sapphire) and a 637 nm laser (Coherent OBIS) through a quartz prim (Tower Optical Co.). For imaging SYTOX Orange-stained DNA, the 488 nm laser power was adjusted to deliver low power (4 mW) at the front face of the prism using a neutral density filter set (Thorlabs). For short DNA-ATTO647N capture experiments, the 637 nm laser power was adjusted to 10 mW. Images were recorded using two electron-multiplying charge-coupled device (EMCCD) cameras (Andor iXon DU897). DNA was extended via continuous buffer flow (0.15 mL min^-1^) in imaging buffer (40 mM Tris-HCl pH 7.5, 25 mM NaCl, 2 mM MgCl_2_, 0.2 mg mL^-1^ BSA, 1 mM DTT, 100 nM SYTOX Orange) supplemented with an oxygen scavenging system (3% D-glucose (w/v), 1 mM Trolox, 1500 units catalase, 250 units glucose oxidase; all from Sigma-Aldrich). Unless indicated, 500 μM NAD^+^ (Spectrum Chemical) was added in the imaging buffer. NIS-Elements software (Nikon) was used to collect the images at a 1-5 s frame rate with 80 ms exposure time. All images were exported as uncompressed TIFF stacks for further analysis in FIJI (NIH) and MATLAB (The MathWorks).

Flowcells were assembled with a 4-mm-wide, 100-μm-high flow channel between a glass coverslip (VWR) and a custom-made quartz microscope slide using two-sided tape (3M). DNA curtains were prepared with 40 μL of liposome stock solution (97.7% DOPC (Avanti #850375P), 2.0% DOPE-mPEG2k (Avanti #880130P), and 0.3% DOPE-biotin (Avanti #870273P) in 960 μL Lipids Buffer (10 mM Tris-HCl, pH 8.0, 100 mM NaCl) incubated in the flowcell for 30 minutes. Then, the flowcell was washed with BSA Buffer (40 mM Tris–HCl, pH 8.0, 2 mM MgCl2, 1 mM DTT, 0.2 mg mL^-1^ BSA) and incubated for 10 minutes. Streptavidin (0.1 mg mL^-1^ diluted in BSA buffer) was injected into the flowcell for another 10 min. Finally, ∼12.5 ng μL^-1^ of DNA substrate was introduced into the flowcell. Subsequently, 100 μL of 1 unit mL^-1^ SfoI restriction enzyme (NEB) was injected to generate blunt-end DNA molecules.

#### DNA substrates for single-molecule imaging

To prepare DNA substrates for microscopy, 125 μg of λ-phage DNA was mixed with two oligos (2 μM oligo Lab07 (/5Phos/AGG TCG CCG CCC/3BioTEG) and 2 μM oligo Lab06 (/5Phos/GGG CGG CGA CCT/3BioTEG) in 1× T4 DNA ligase reaction buffer (NEB B0202S) and heated to 70°C for 15 min followed by gradual cooling to 15°C for 2 hours. One oligo will be annealed with the overhand located at the left cohesive end of DNA, and the other oligo will be annealed with the overhang at the right cohesive end. After the oligomer hybridization, 2 μL of T4 DNA ligase (NEB M0202S) was added to the mixture and incubated overnight at room temperature to seal nicks on DNA. The ligase was inactivated with 2 M NaCl, and the reaction was resolved over a custom-packed S-1000 gel filtration column (GE) to remove excess oligonucleotides and proteins.

For the preparation of 60 bp short DNA labeled with ATTO647N, Oligo1 (/5BioTEG/ACG AAG TCT TAT GGC AAA ACC GAT GGA CTA TGT TTC GGG TAG CAC CAG AAG TCT ATA ACA) and ATTO647N-tagged Oligo2 (5TGT TAT AGA CTT CTG GTG CTA CCC GAA ACA TAG TCC ATC GGT TTT GCC ATA AGA CTT CGT /3ATTO647N/) were purchased from IDT and annealed by combining 20 μM Oligo1 with 20 μM Oligo2. The annealing process involved heating to 75°C for 10 minutes, followed by a gradual cooling to 22°C over a period of 1 hour in a thermal cycler.

#### Ligation assay

PARylation reactions were set up as described above before with Cy5-labeled triplex DNA and ATP added to allow ligation at 37°C for the indicated period of time before quenching the reaction with 2x quenching buffer (1x TBE, 12% Ficoll 400, 7 M urea) to a final concentration of 1x. Reactions were then incubated at 95°C for 5 min to denature the DNA. Samples were loaded on a denaturing 7 M urea gel and resolved for 30 min 180 V. Gels were imaged using the Cy5 filter on a BioRad GelDoc EZ system. Ligation efficiency analysis was performed on Fiji by obtaining the ratio of the ligated and unligated product.

---

**Supplemental Table 1.** Phase separation predictions for proteins investigated in this study using PScore, CatGRANULE, and PLAAC prediction algorithms. PScore (99) uses pi-pi interactions, PLAAC (100) uses prion-like domain features, and catGRANULE (101) uses similarities to sequence features found in RNA granule components. Values reflect percentiles of scores within the human proteome, with 100 percent indicating the highest likelihood for phase separation (102).

| Name | UniProt ID | PScore | CatGRANULE | PLAAC |
| --- | --- | --- | --- | --- |
| **PARP1** | P09874_HUMAN | 62.36 | 96.31 | 54.59 |
| **XRCC1** | P18887_HUMAN | 65.47 | 86.39 | 73.13 |
| **LigIII** | P49916_HUMAN | 66.35 | 86.10 | 58.98 |
| **Polβ** | P06746_HUMAN | 18.36 | 82.30 | 9.743 |

**Supplemental Table 2.**
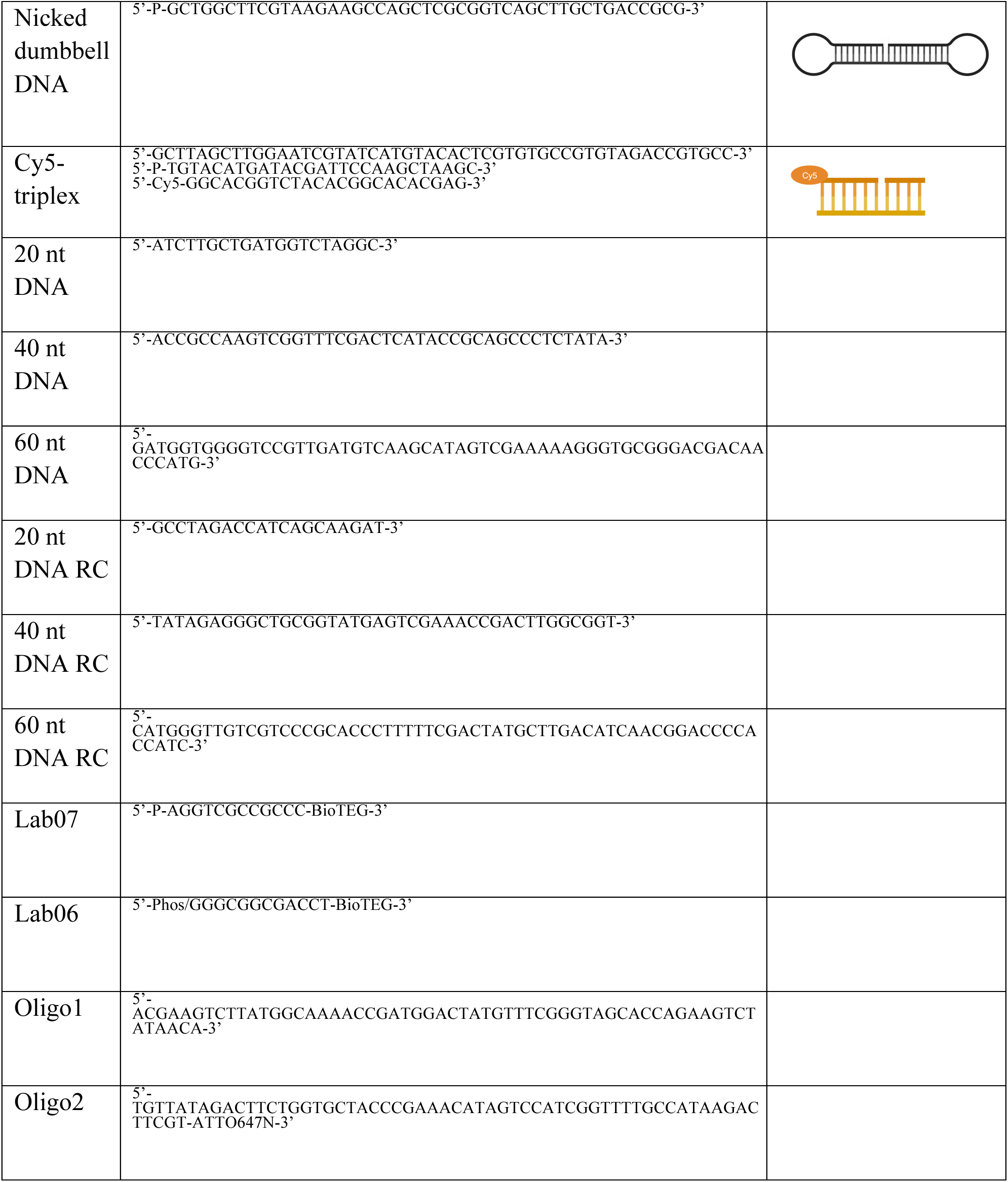
Sequences of oligonucleotides used in this study.

## ACKNOWLEDGEMENTS

We would like to thank members of the Forman-Kay, H. Lee, and Ditlev labs (Biochemistry, University of Toronto) for stimulating discussions and critical reading of the manuscript. We thank Fiona Cui for assistance with bioinformatic analysis. C.C.S. and M.L.N. are funded by a Canada Graduate Scholarship-Doctoral from the Natural Sciences and Engineering Research Council (NSERC). M.T. is funded by a NSERC Vanier Canada Graduate Scholarship. H.Z. and I.J.F. are funded by NCI grant 5P01CA092584 and the Welch Foundation (F-1808). T.R.A. is funded by a Banting Postdoctoral Fellowship from the Canadian Institutes of Health Research (CIHR). This work was funded from NSERC RGPIN-2019-070 to H.O.L and CIHR FDN-148375 to J.D.F.-K., and the Canada Research Chair program to both J.D.F.-K. and H.O.L.

## CONFLICTS OF INTEREST

The authors declare no conflicts of interest.

## AUTHOR CONTRIBUTIONS

C C.S., G.M., M.T., M.L.N, J.D.F.-K. and H.O.L. contributed to the conceptualization, investigation and analysis of the results, and writing of the manuscript. H.Z. and I.J.F. contributed to the conceptualization, investigation and analysis of the results, and writing of the single-molecule microscopy-related experiments. T.R.A. provided caspase-3 protein and contributed to conceptualization of caspase-3-related experiments. M.D. and A.L. contributed to the conceptualization of the PAR chain-related experiments and provided purified PAR chains, performed PAR concentration and chain length experiments, analysis, and writing of the PAR chain-related experimental methods.

---

**Supplemental Figure 1.**
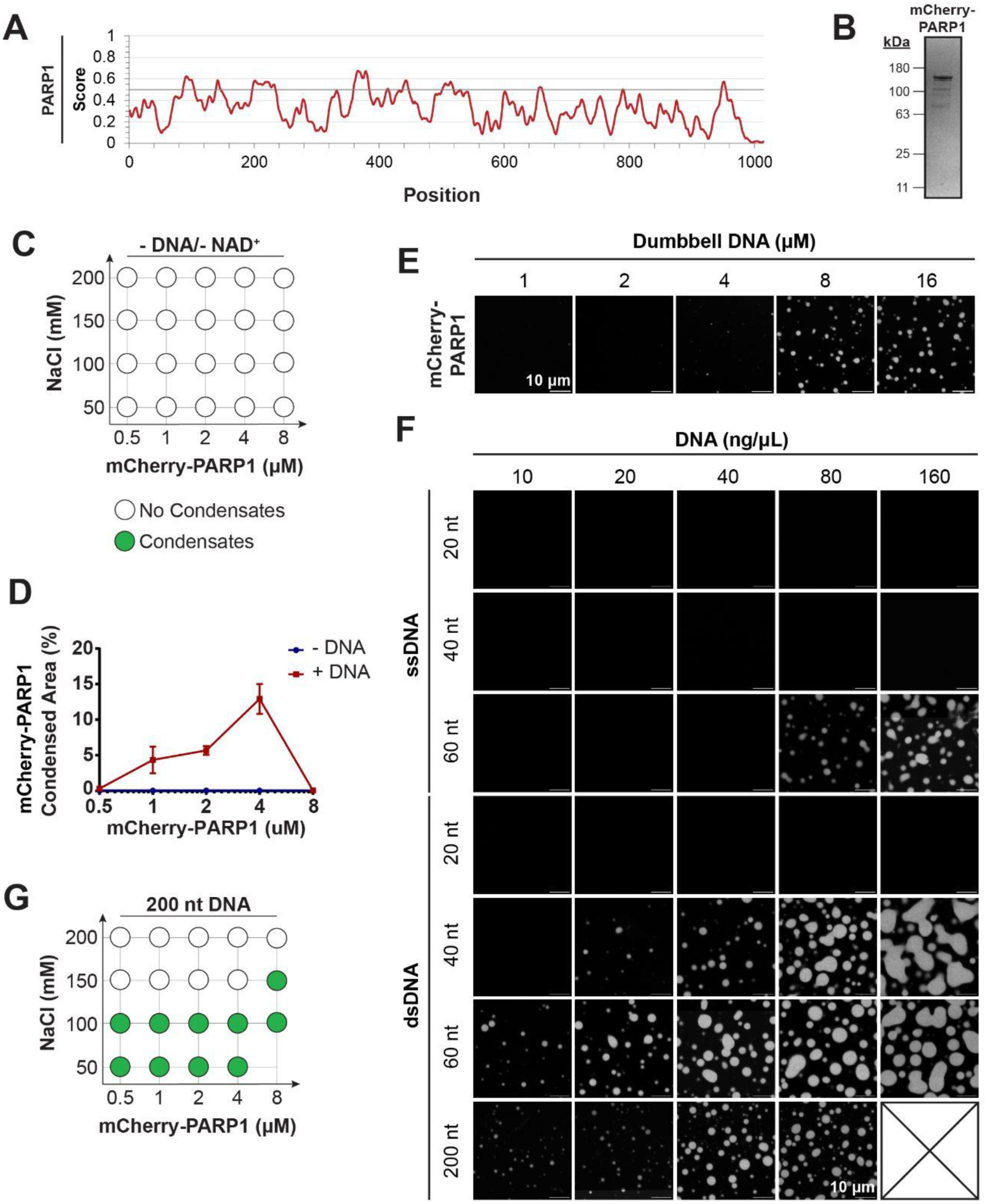
PARP1 forms condensates in a DNA-dependent manner. **A**. IUPRED3 disordered region plots for PARP1 (https://iupred.elte.hu/plot). **B.** SDS-PAGE of purified recombinant mCherry-PARP1; the gel was stained with Coomassie Blue. **C.** Condensate formation by mCherry-PARP1 at the indicated protein and salt concentrations in PARylation Buffer (20 mM Tris pH 7.5, 50 mM NaCl, 7.5 mM MgCl_2_ and 1 mM DTT) in the absence of DNA and NAD^+^. **D.** Quantifications of the percent of surface area covered by mCherry-PARP1 condensates in Fig. 1C. Error bars indicate the standard error of the mean (SEM). **E.** Fluorescence micrographs of 4 µM mCherry-PARP1 in PARylation Buffer with the indicated concentration of Dumbbell DNA without NAD^+^. **F.** Fluorescence micrographs of 4 µM mCherry-PARP1 in PARylation Buffer with the indicated concentration of single-stranded DNA (ssDNA) and double-stranded DNA (dsDNA) of 20, 40, 60 and 200 nucleotides in length without NAD^+^. **G.** Condensate formation by 4 µM Cherry-PARP1 with the indicated concentration of 200-bp DNA in PARylation Buffer in the absence or presence of NAD^+^. All experiments were repeated in at least 3 independent replicates.

**Supplemental Figure 2.**
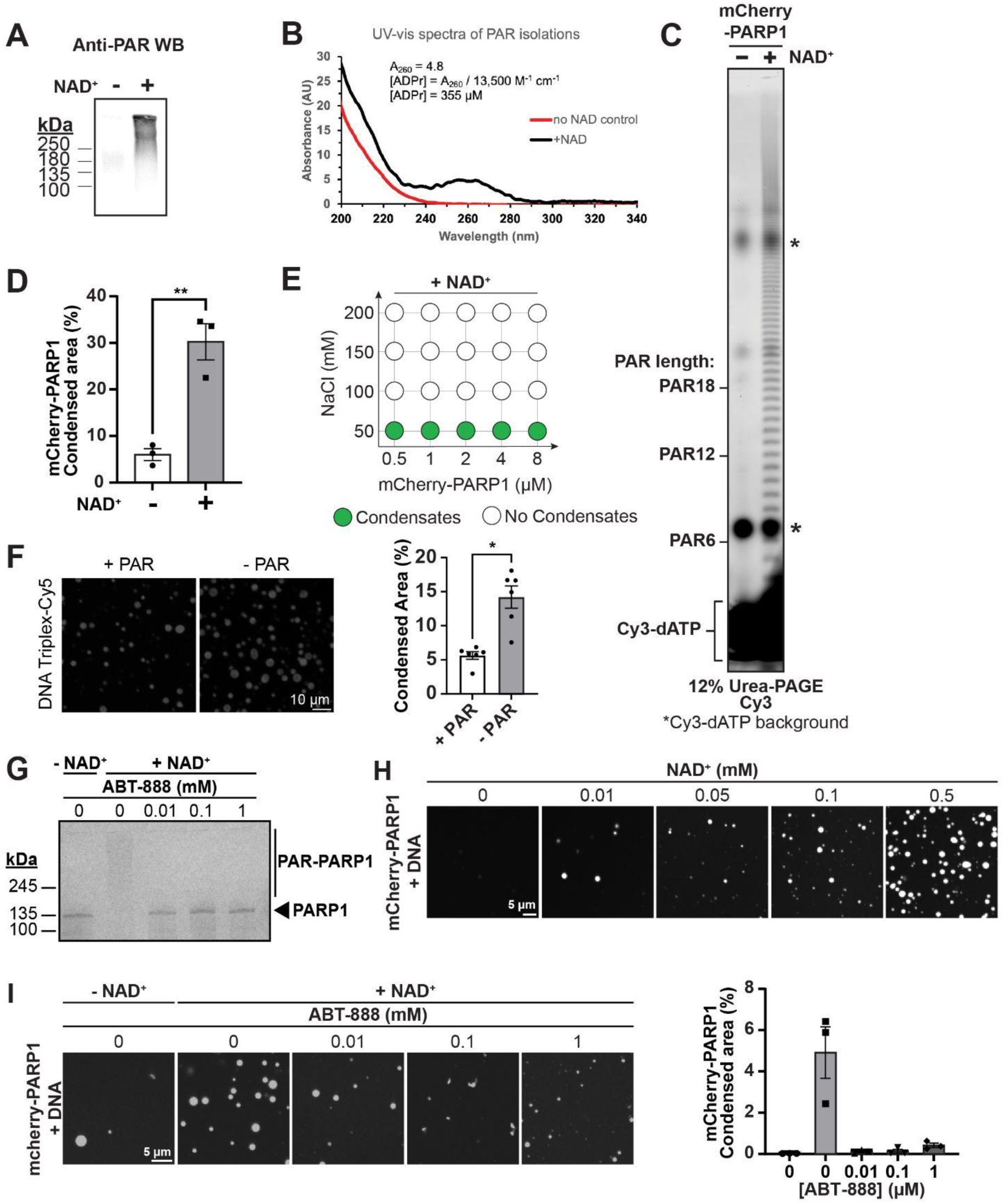
AutoPARylation enhances formation and internal dynamics of PARP1 condensates. **A.** Western blot (anti-PAR) depicting autoPARylation of PARP1 from reactions of 1 µM untagged PARP1 incubated with 0.3 µM dumbbell DNA in PARylation Buffer (20 mM Tris pH 7.5, 50 mM NaCl, 7.5 mM MgCl_2_ and 1 mM DTT) for 15 min with and without 0.5 mM NAD^+^. **B.** UV-Vis spectra of mCherry-PARP1 with 0.3 µM dumbbell DNA in PARylation Buffer for 15 minutes with and without 0.5 mM NAD^+^. ADP-ribose concentration is calculated with the equation depicted and is estimated to be around 355 µM. **C.** Urea-PAGE of PAR extracted from mCherry-PARP1 in PARylation reactions with and without 0.5 mM NAD^+^, labeled with Cy3-dATP. **D.** Quantifications of the percent of the surface area covered by mCherry-PARP1 condensates at 4 µM mCherry-PARP1 and 1.2 µM triplex DNA with and without NAD^+^ represented in Figure 1B. Error bars indicate the standard error of the mean (SEM). **E.** Condensate formation by mCherry-PARP1 at the indicated protein and salt concentrations in an autoPARylation reaction in PARylation Buffer with 0.5 mM NAD^+^. **F.** Left: Fluorescence micrographs of 2 µM Cy5-Triplex DNA with 4 µM mCherry-PARP1 and 0.5 µM PAR16 chains in PARylation Buffer. Right: Quantifications of the surface area covered by Cy5-triplex DNA condensates. Error bars indicate the SEM. The Cy5-triplex DNA signal is displayed on different ranges to be able to visualize the low signal upon PAR addition. **G.** Gel-based autoPARylation assay performed with 1 µM mCherry-PARP1 in PARylation Buffer with with 0.3 µM triplex DNA, with or without 0.5 mM NAD^+^, and increasing concentrations of ABT-888 (0 mM, 0.01 mM, 0.1 mM, and 1 mM). **H.** Fluorescence micrographs of 1 µM mCherry-PARP1 in PARylation Buffer with 0.3 µM triplex DNA indicated concentrations of NAD^+^. **I.** Fluorescence micrographs of 1 µM mCherry-PARP1 in PARylation Buffer with 0.3 µM triplex DNA, with or without 0.5 mM NAD^+^, and indicated concentrations of ABT-888. All experiments were repeated in at least 3 independent replicates unless specified otherwise. p-values are obtained from a student t-test: ** p < 0.01, * p < 0.05.

**Supplemental Figure 3.**
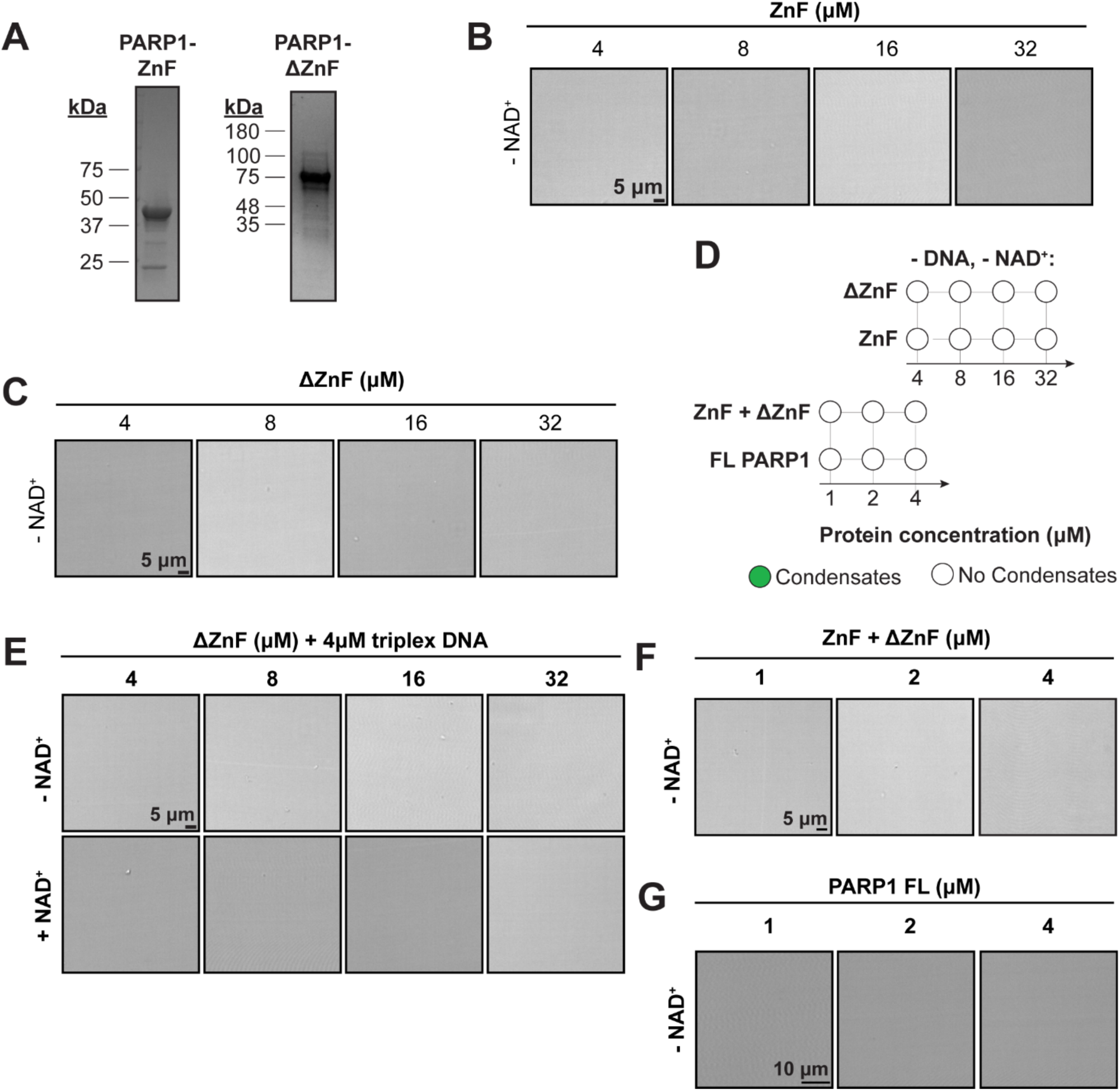
The ZnF region is important for PARP1 condensation. **A**. Coomassie-stained SDS-PAGE gels of purified recombinant PARP1 truncation constructs. **B**. DIC micrographs of ZnF PARP1 protein without triplex DNA in PARylation Buffer (20 mM Tris pH 7.5, 50 mM NaCl, 7.5 mM MgCl_2_, 1 mM DTT). **C**. DIC micrographs of ΔZnF PARP1 protein without triplex DNA in PARylation Buffer. **D**. Condensate formation by truncated and full-length PARP1 proteins at the indicated concentrations in PARylation Buffer. **E**. DIC micrographs of ΔZnF PARP1 protein fragments with 4 µM triplex DNA, and with 0.5 mM NAD^+^ in PARylation Buffer. **F**. DIC micrographs of ΔZnF PARP1 protein without triplex DNA in PARylation Buffer. **G.** DIC micrographs of full-length PARP1 protein fragments without triplex DNA in PARylation Buffer. All experiments were repeated in at least 3 independent replicates unless specified otherwise.

**Supplemental Figure 4:**
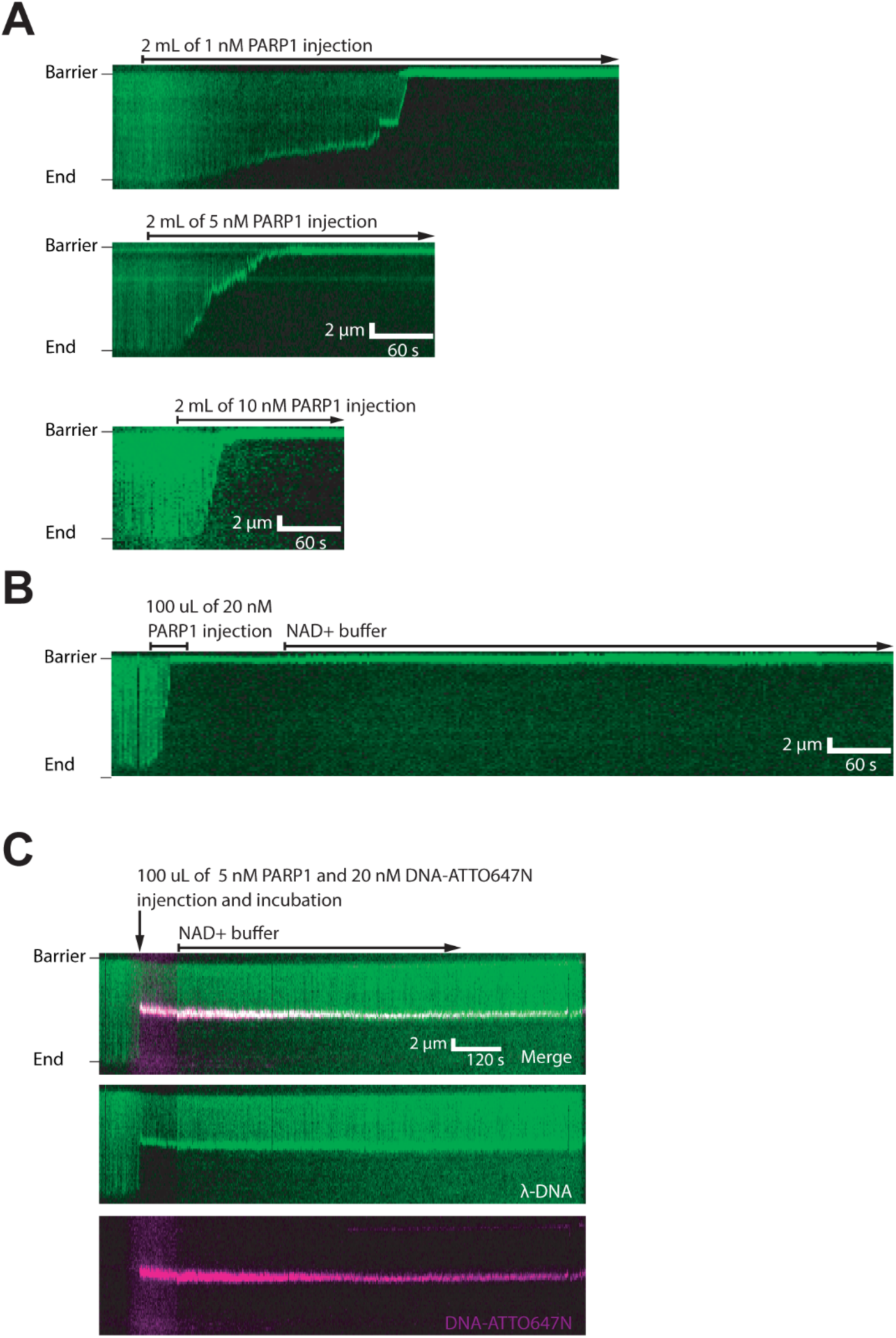
PARP1 condensates bridge broken DNA ends. **A.** Kymographs showing the compaction of DNA (green) induced by the injection of 1 nM, 5 nM, and 10 nM PARP1, respectively. **B.** Kymographs illustrating DNA compaction by 100 μL of 20 nM PARP1, followed by a buffer switch to the imaging buffer containing 500 μM NAD^+^. **C.** Kymographs showing the capture of 60 bp DNA-ATTO647N induced by PARP1. A mixture of 100 μL, containing 5 nM PARP1 and 20 nM DNA-ATTO647N, was injected into the flowcell and incubated with tethered λ-DNA (in green) with flow turned off. After a 5-minute incubation, imaging acquisition resumed, revealing the capture of DNA-ATTO647N (in magenta) on the long DNA (green). Two minutes later, the imaging buffer was supplemented with 500 μM NAD^+^.

**Supplemental Figure 5.**
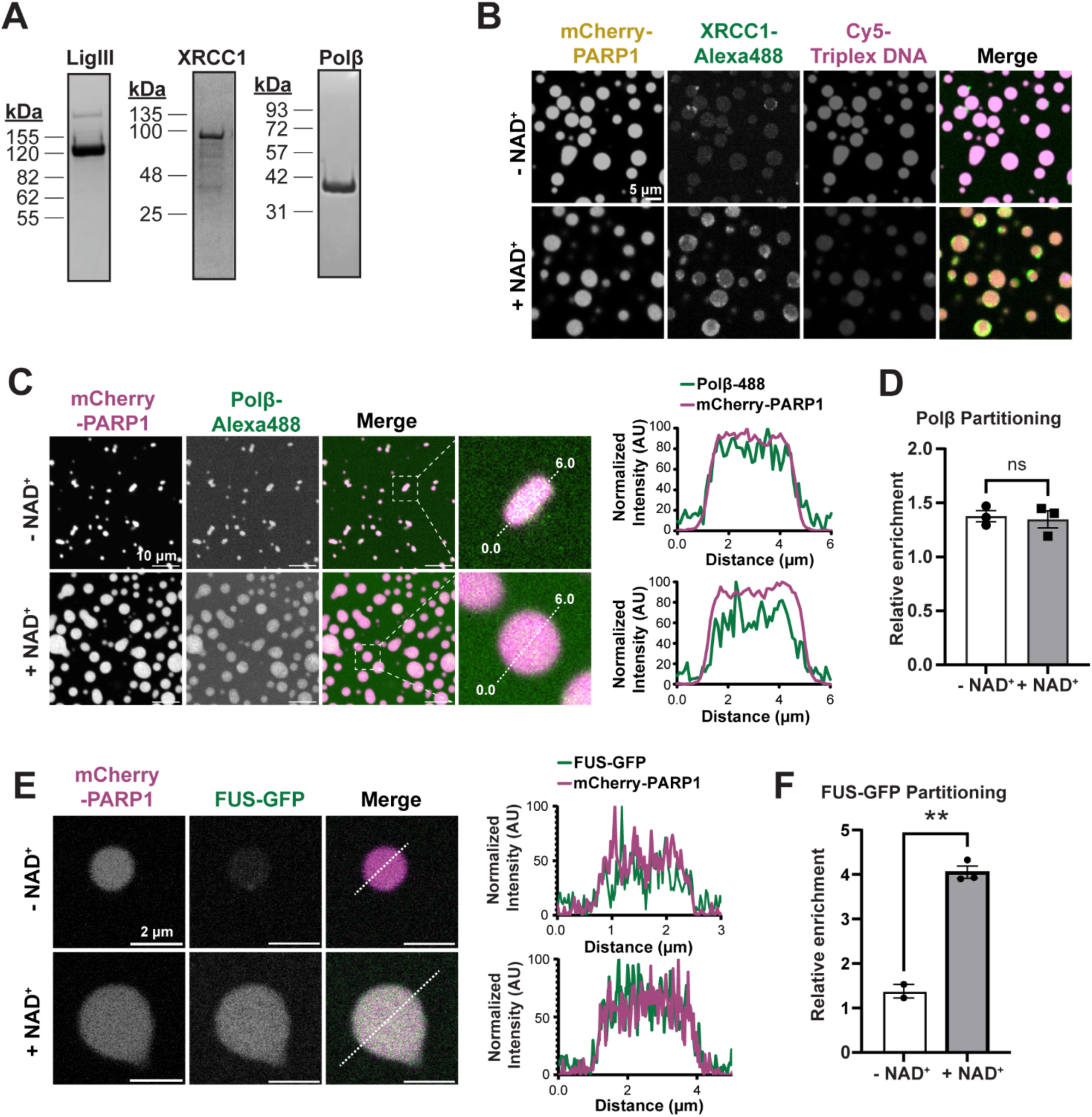
SSBR proteins partition together in condensates that enrich DNA. **A.** Coomassie-stained SDS-PAGE gels of purified recombinant LigIII, XRCC1, Polβ and FUS-GFP. **B.** Fluorescence micrographs of 4 µM mCherry-PARP1, 4 µM triplex DNA (10% Cy5-labeled triplex DNA) and 1 µM XRCC1 (10% AlexaFluor488-labeled XRCC1) in PARylation Buffer (20 mM Tris pH 7.5, 50 mM NaCl, 7.5 mM MgCl_2_, 1 mM DTT) with and without NAD^+^. The Cy5-triplex signal is displayed on different ranges to be able to visualize the low signal upon NAD^+^ addition. Quantifications of the XRCC1 partitioning are shown in Fig. 5B. **C.** Fluorescence micrographs of 4 µM mCherry-PARP1, 8 µM dumbbell DNA and 1 µM Polβ (10% AlexaFluor488-labeled Polβ) in PARylation Buffer with and without NAD^+^. The fluorescence intensity measured across the diameter of a representative (dashed white line) condensate is plotted in the center. **D.** Quantification of the images in C. The enrichment of the Polβ within condensates is calculated by the mean fluorescence intensity in the condensed phase relative to that of the dilute phase. **E.** Fluorescence micrographs of 4 µM mCherry-PARP1, 1 µM triplex DNA, and 1 µM FUS-GFP in PARylation Buffer with and without NAD^+^. The fluorescence intensity measured across the diameter of a representative condensate (white dashed line) is plotted. **F.** Quantifications of the images in E. The enrichment of the FUS-GFP within condensates as calculated by the mean fluorescence intensity in the condensed phase relative to that of the dilute phase is plotted. All experiments were repeated in at least 3 independent replicates unless specified otherwise. p-values are obtained from a student t-test: ** p < 0.01, n.s.: not significant.

## Notes

### Competing Interest Statement

The authors have declared no competing interest.

